# Rescue of alveolar wall liquid secretion blocks fatal lung injury by influenza-staphylococcal coinfection

**DOI:** 10.1101/2022.07.07.499169

**Authors:** Stephanie Tang, Ana Cassandra De Jesus, Deebly Chavez, Sayahi Suthakaran, Sarah K. L. Moore, Raveen Rathnasinghe, Michael Schotsaert, Jahar Bhattacharya, Jaime L. Hook

## Abstract

Secondary lung infection by inhaled *Staphylococcus aureus* (SA) is a common and lethal event in individuals infected with influenza A virus (IAV). It is unclear how IAV disrupts host defense to promote SA infection in lung alveoli, where fatal lung injury occurs. We addressed this issue using the first real-time determinations of alveolar responses to IAV in live, intact, perfused lungs. Our findings show IAV infection blocked defensive alveolar wall liquid (AWL) secretion and induced airspace liquid absorption, thereby reversing normal alveolar liquid dynamics and inhibiting alveolar clearance of inhaled bacteria. Loss of AWL secretion resulted from dephosphorylation, hence inhibition of the cystic fibrosis transmembrane conductance regulator (CFTR) ion channel in alveolar epithelium, and airspace liquid absorption was caused by alveolar epithelial stimulation of the epithelial Na^+^ channel (ENaC). Loss of AWL secretion promoted alveolar stabilization by SA and alveolar damage by the secreted SA toxin, alpha hemolysin, but rescue of AWL secretion protected against fatal SA-induced lung injury in IAV-infected mice. These findings reveal a central role for AWL secretion in alveolar defense against inhaled bacteria and identify AWL inhibition as a critical mechanism of IAV lung pathogenesis. AWL rescue may represent a new therapeutic approach for IAV-SA coinfection.

**GRAPHICAL ABSTRACT:** 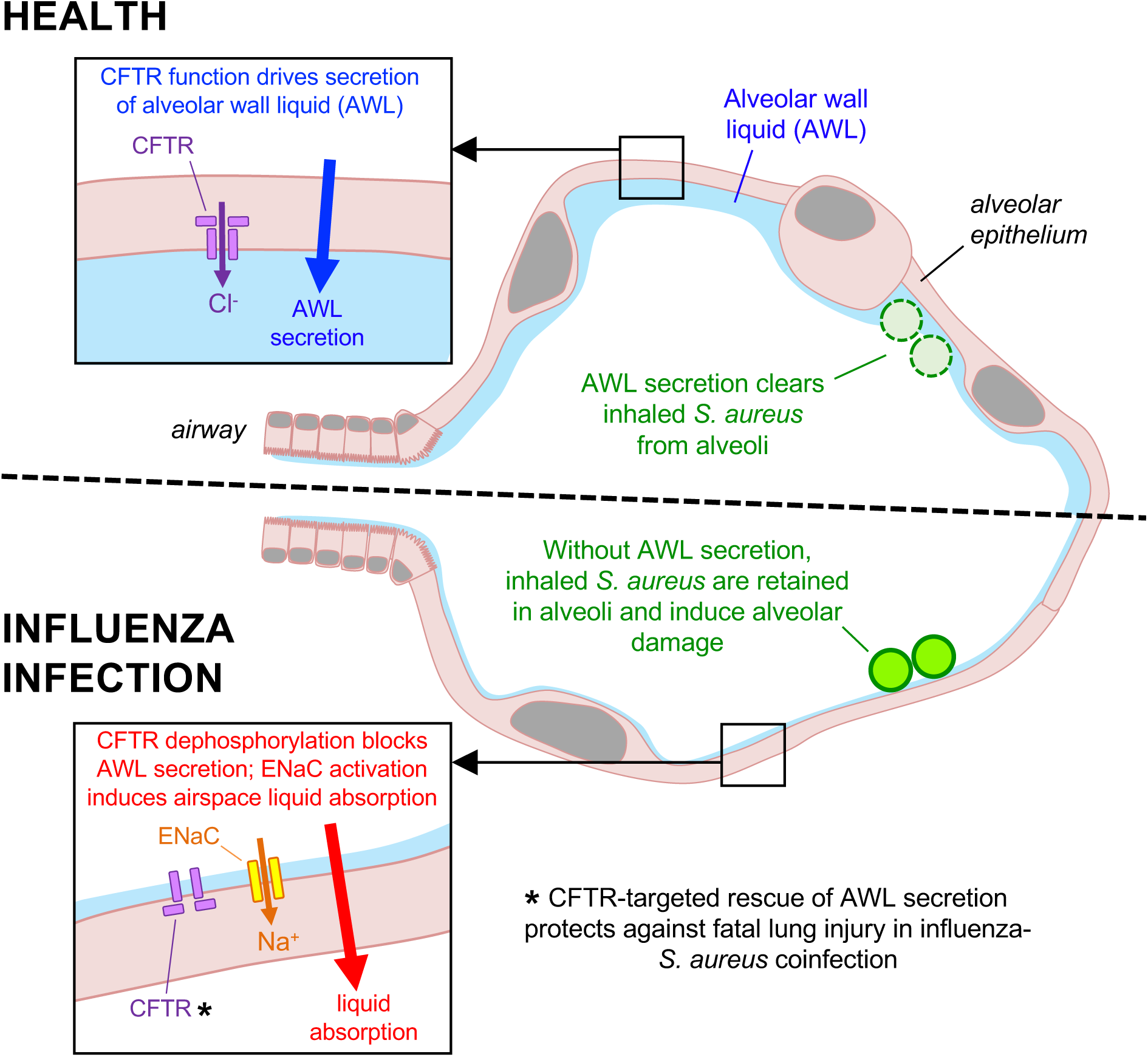

## INTRODUCTION

Lung infection is the fourth leading cause of global mortality (1), and the majority of severe lung infections are caused by respiratory viruses (2, 3). Among virus-infected individuals, the highest rates of death occur in those who acquire secondary lung infection by inhaled bacteria (4–9). Factors that account for the virulence of viral-bacterial coinfection remain unclear. Since coinfection pathogenesis centers on acute lung injury (10, 11), a disease of lung alveoli (12, 13), coinfection virulence might result from virus-induced alveolar responses that render alveoli susceptible to bacterial colonization and toxicity. However, there is little understanding as to how alveoli respond to respiratory viruses, including those responses that promote secondary bacterial infection. Here, we consider these issues in the context of lung coinfection by influenza (IAV) and *Staphylococcus aureus* (SA), a common and lethal (4–7) pathogen combination for which current therapy is insufficiently effective (14) and increasingly hindered by pathogen drug resistance (15, 16).

Known mechanisms by which IAV promotes secondary SA infection and SA-induced lung injury implicate the lung airways, microvessels, and immune system. Inhaled SA encounter IAV-induced airway epithelial changes that promote bacterial adherence, including cell loss that exposes bacterial attachment sites (17) and impaired mucociliary velocity that hampers bacterial clearance (18). Lung persistence of SA is enhanced by IAV-induced immune cell dysfunction that impairs bacterial uptake and increases bacterial survival (19, 20). Subsequently, SA cause acute lung injury by inducing inflammatory cell recruitment (21), innate immune cell necrosis (22), and microvascular endothelial barrier dysfunction (23). Although these mechanisms address major aspects of coinfection pathogenesis, they do not account for the initiation of SA infection or SA-induced tissue damage in alveoli. This issue is important because alveoli comprise more than 95% of the lung surface area (24, 25) and critically determine the alveolar fluid barrier properties that maintain lung function (12, 13).

We previously used real-time optical imaging of live, intact lungs to define alveolar mechanisms by which inhaled SA alone initiate alveolar infection and induce alveolar damage (26). Our published findings show bacteria of the SA strain, USA300 inhaled while in the exponential growth phase rapidly form stable microaggregates in structural niches of alveolar walls, where SA secretion of the epithelial-damaging toxin, alpha hemolysin (Hla), leads to alveolar epithelial barrier dysfunction, alveolar airspace edema, and fatal acute lung injury (26). The ability of USA300 SA to stabilize in alveolar niches is critical to its lung virulence, since bacteria that fail to stabilize in alveoli do not induce alveolar damage (26). Thus, our findings raise the question as to how non-USA300 SA, which may be less apt to stabilize in alveoli, succeed at causing clinical lung injury after IAV lung infection (27). Our hypothesis is IAV lung infection promotes alveolar stabilization of inhaled bacteria by disrupting endogenous mechanisms of alveolar defense. Since IAV inhibits the cystic fibrosis transmembrane conductance regulator (CFTR) ion channel in vitro (28–30), IAV may block CFTR-dependent secretion of alveolar wall liquid (AWL), a homeostatic epithelial function that clears inhaled particles from alveoli (31). We tested this hypothesis by carrying out the first determinations of alveolar responses to IAV lung infection in live, intact, perfused lungs.

## RESULTS

### IAV lung infection blocks AWL secretion and induces alveolar liquid absorption

We used real-time confocal microscopy to view live alveoli of intact, blood-perfused lungs of mice that were untreated or intranasally instilled with IAV at 24 h prior to lung excision for imaging. To evaluate the effect of IAV lung infection on AWL secretion, we first determined alveolar epithelial viability and fluid barrier function in IAV-infected lungs, respectively, by microinstillation of calcein dye into alveolar airspaces and addition of fluorophore-labeled dextran (20 kD) to the lung perfusate solution. In IAV-infected lungs, cytosolic calcein fluorescence (Figure 1A) indicates the alveolar epithelium was viable, and confinement of dextran fluorescence to alveolar microvessels (Figure 1, B and C) indicates alveolar fluid barrier function was intact.

**Figure 1.**
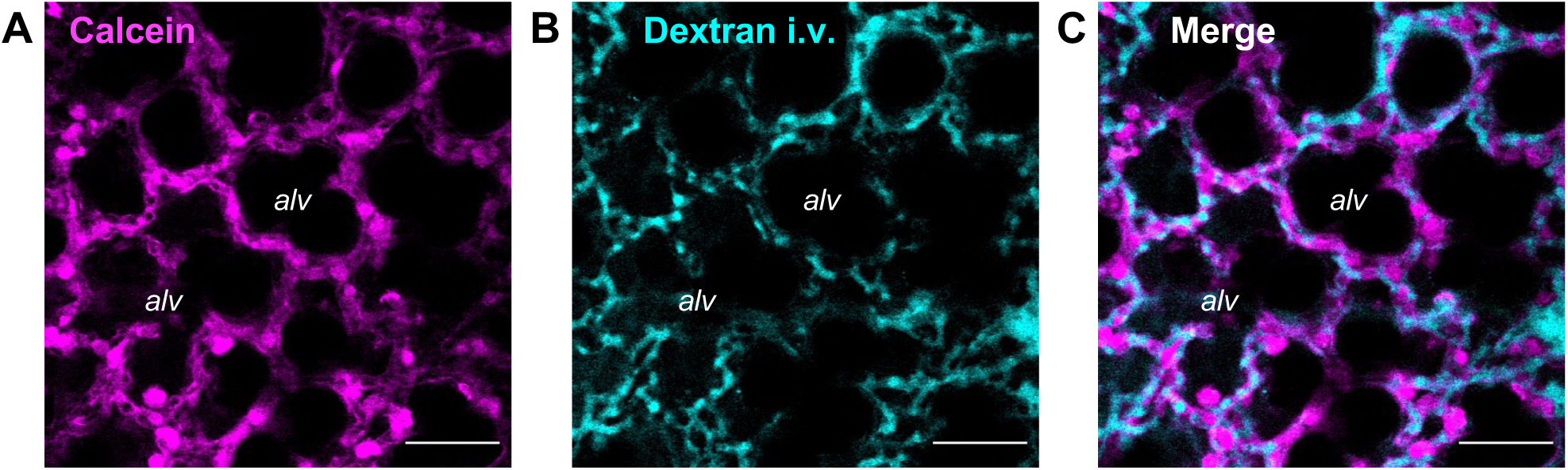
Alveolar epithelial viability and barrier function in IAV-infected lungs. (**A-C**) Confocal images show epithelial fluorescence of calcein (*magenta*) and intravascular (*i.v.*) fluorescence of tetramethylrhodamine-labeled dextran (70 kD; 10 mg/mL; *cyan*) in live alveoli of intact, blood-perfused mouse lungs. Lungs were excised for imaging at 24 h after intranasal IAV instillation. Calcein-AM was microinstilled in alveoli by alveolar micropuncture, and dextran was added to the lung perfusate solution. Note, dextran fluorescence is absent from alveolar airspaces (*alv*) in the merged image (C). Scale bars: 50 µm. Replicated in lungs of 3 mice.

To visualize the AWL, we used an established approach (26, 31) in which we perfused the lungs with non-fluorescent blood-buffer solution, then micropunctured a single alveolus under bright-field microscopy to instill alveolar airspaces with a 2-s microinfusion of fluorophore-labeled dextran (70 kD) in aqueous solution. The microinfusion spread to airspaces of at least 20 neighboring alveoli, as evident by transient loss of optical discrimination between alveolar walls and airspaces. Return of optical discrimination occurred within seconds of each microinstillation, indicating free fluid rapidly drained from alveoli and reestablished the air-filled alveolar lumens (32). In line with the findings of others (31), confocal imaging revealed dextran fluorescence in airspaces as a juxta-epithelial layer that accumulated at alveolar niches (Figure 2A, arrowheads), curved regions of alveolar walls where septa converge (26). Airspace washout by alveolar microinfusion of non-fluorescent buffer abolished the fluorescence (data not shown), indicating dextran was restricted to airspaces and not taken up by the alveolar wall.

**Figure 2.**
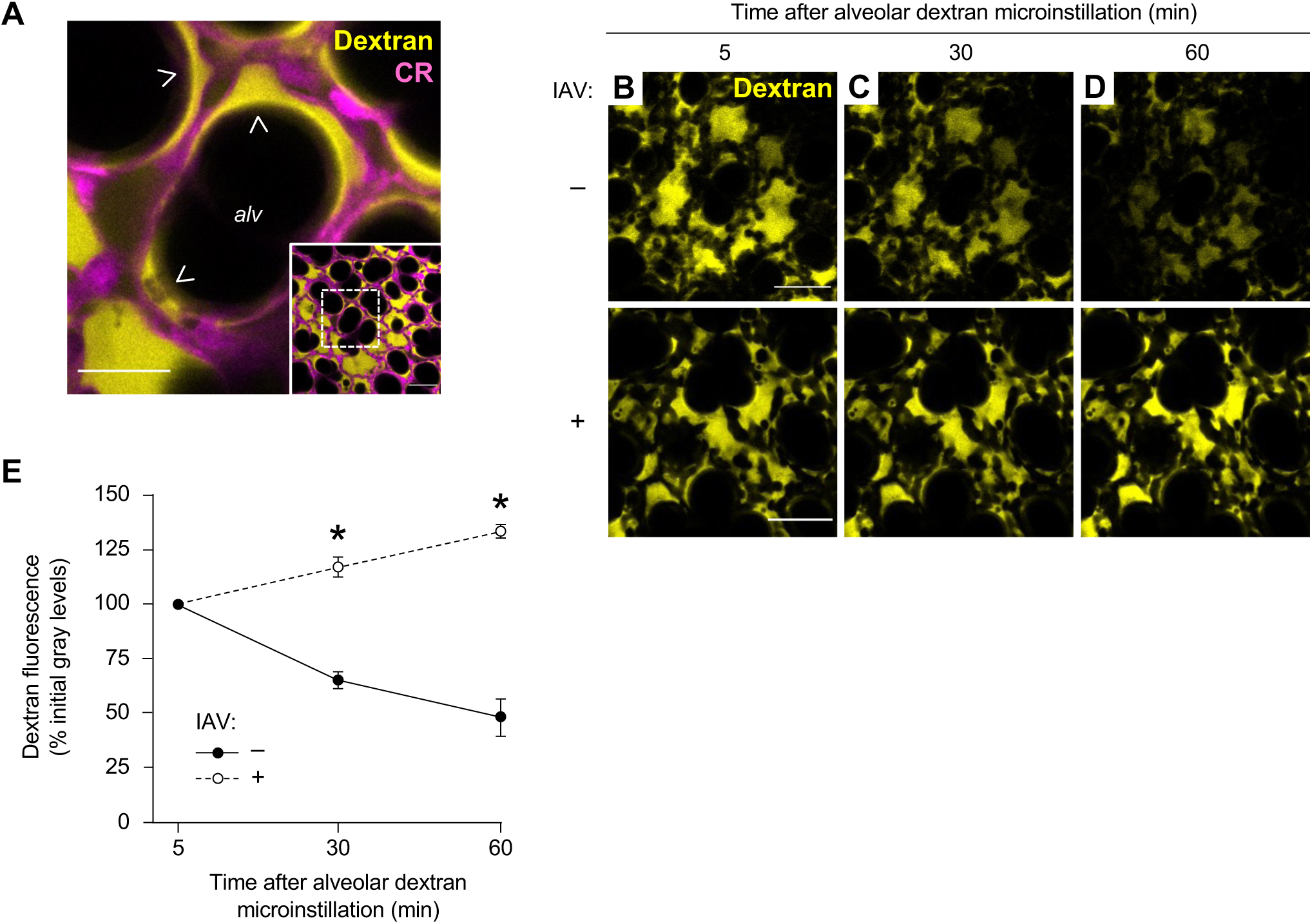
IAV lung infection disrupts AWL secretion in live alveoli. (**A**) Low (*inset*) and high power confocal images show fluorescence of tetramethylrhodamine (TRITC)-conjugated dextran (70 kD; 10 mg/mL; *yellow*) in live alveoli (*magenta*) at 5 min after alveolar dextran microinstillation. Note, dextran formed a thin layer against alveolar walls and pooled in structural alveolar niches (*arrowheads*). *CR*, calcein red-orange; *alv,* alveolar airspace. Scale bars: 50 (*inset*) and 20 µm. Images replicated in 40 mice. (**B-E**) Confocal images (B-D) and group data (E) show time-dependent change of alveolar dextran fluorescence in airspaces of live alveoli in lungs excised from mice that were untreated (B-D, *top row* and E, *closed circles*; *n*=4 mice) or intranasally instilled with IAV at 24 h prior to imaging (B-D, *bottom row* and E, *open circles*; *n*=4 mice). Fluorescence of alveolar walls is not shown. Group data (E) represent mean ± SEM. For each mouse, mean dextran fluorescence was quantified at each of the 3 indicated time points in an imaging field containing at least 30 lung alveoli. \**P* < 0.05 versus *closed circles* by two-tailed *t* test. Scale bars: 50 µm.

Calibration experiments in glass micropipettes show dextran fluorescence varied with dextran concentration (Supplemental Figure 1, A and B) and was unchanged after repeated imaging (data not shown). However, dextran fluorescence decreased over time in alveolar airspaces of unchallenged lungs (Figure 2, B-D, top row and Figure 2E, closed circles), confirming published findings (31) and indicating the dextran was progressively diluted by addition of non-fluorescent liquid. To determine whether the dilution resulted from airspace accumulation of airway liquid, we inferred time-dependent change of airspace dextran volume from quantifications of dextran pool width at alveolar niches in high-power images at a specific distance below the pleura (Figure 3, A-E). Our findings align with published data (31) and show airspace and dextran pool widths were steady in alveoli of unchallenged lungs (Figure 3E, first and second bars, respectively). These findings show there was no inflow of liquid from the airways during the period of dextran fluorescence loss. Hence, we interpret airspace dextran dilution resulted from alveolar liquid secretion, confirming reports (31) that the alveolar epithelium continuously secretes AWL into alveolar airspaces under baseline conditions.

**Figure 3.**
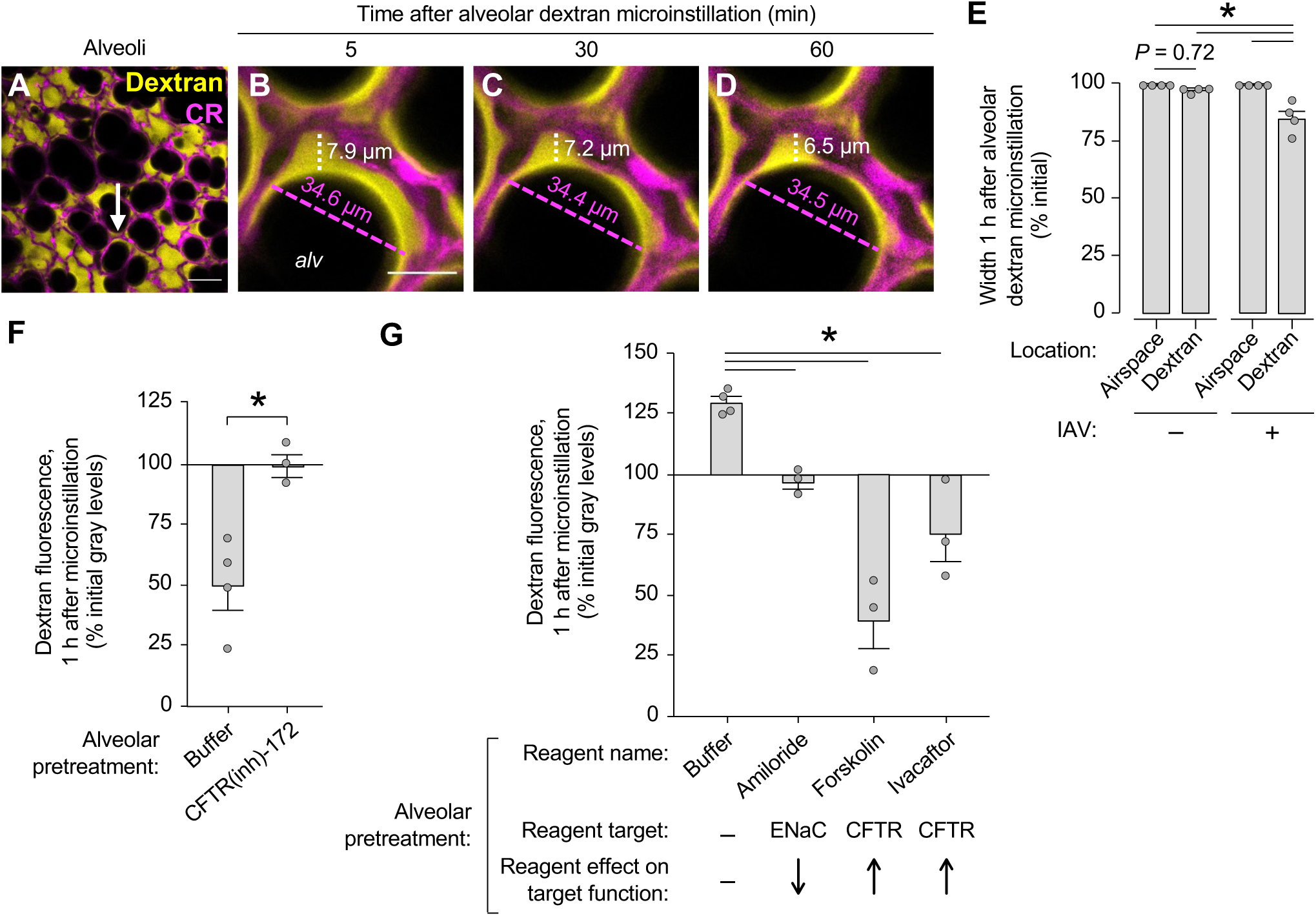
IAV lung infection induces airspace liquid absorption in live alveoli. (**A-E**) Confocal images (A-D) and group data (E) show time-dependent change of fluorescence of microinstilled TRITC-conjugated dextran (70 kD; 10 mg/mL; *yellow*) in live alveoli (*magenta*). Imaged lungs were excised from mice that were untreated ((-), shown in E only) or intranasally instilled with IAV ((+), A-E) at 24 h prior to excision. High power confocal views (B-D) of the structural alveolar niche indicated by the *arrow* (A) demonstrate, in IAV-infected lungs, time-dependent decrease of dextran pool width (*white dashed line* and *white text*) but not airspace width (*magenta dashed lines* and *magenta text*). For group data (E), circles indicate *n* and each represent one mouse in which widths were quantified at 10 random locations in an imaging field containing at least 30 alveoli. Bars: mean ± SEM; \**P* < 0.05 as indicated by ANOVA with post hoc Tukey testing. *CR,* calcein red-orange; *alv,* alveolar airspace. Scale bars: 50 (A) and 15 (B) µm. (**F-G**) Group data quantify change of TRITC-dextran fluorescence in alveolar airspaces of live, intact lungs. Mice were untreated (F) or intranasally instilled with IAV (G) at 24 h before lung excision for imaging. Alveolar epithelium was pretreated with alveolar microinstillation of the indicated reagents dissolved in HEPES-buffered solution or buffer alone, as indicated, before alveolar dextran microinstillation. Circles indicate *n* and each represent one mouse in which mean dextran fluorescence change was quantified in an imaging field of at least 30 alveoli. Bars: mean ± SEM; \**P* < 0.05 by two-tailed *t* test (F) or ANOVA with post hoc Tukey testing (G).

By contrast, in alveoli of IAV-infected lungs, we observed time-dependent gain of airspace dextran fluorescence (Figure 2, B-D, bottom row and Figure 2E, open circles), indicating the dextran concentration progressively increased. At the same time, dextran pool width progressively decreased whereas airspace width was unchanged (Figure 3, A-D, white and pink dashed lines and Figure 3E, third and fourth bars). These findings show airspace liquid volume decreased during the period of dextran fluorescence gain, indicating IAV lung infection induced airspace liquid absorption and suggesting IAV inhibited AWL secretion.

To determine mechanisms, we repeated the dextran microinstillation experiments after alveolar epithelial exposure to pharmacologic ion channel activators and inhibitors. First, we confirmed AWL secretion under baseline conditions depends on function of the alveolar epithelial CFTR protein (31) by blocking airspace dextran fluorescence loss with the CFTR inhibitor, CFTRinh-172 (Figure 3F). Second, since lung liquid absorption is determined by activity of the amiloride-sensitive epithelial Na^+^ channel (ENaC) (33, 34), and ENaC is expressed by the alveolar epithelium (35, 36) and drives alveolar epithelial Na^+^ transport (36, 37), we evaluated the role of ENaC in IAV-induced alveolar airspace liquid absorption. Alveolar pretreatment with the ENaC inhibitor, amiloride abolished the dextran fluorescence increase in alveoli of IAV-infected lungs (Figure 3G, second bar), indicating IAV-induced airspace liquid absorption resulted from ENaC stimulation in the alveolar epithelium. Third, alveolar pretreatment with the CFTR activator, forskolin or the CFTR potentiator, ivacaftor, which potentiates channel activity of murine CFTR (38), each induced dextran fluorescence loss in IAV-infected lungs

(Figure 3G, third and fourth bars, respectively). Taking these findings together, we conclude IAV lung infection had a dual effect on the alveolar epithelium characterized by CFTR inhibition and ENaC activation. The result was loss of AWL secretion and stimulation of airspace liquid absorption, leading to net liquid absorption in alveoli of IAV-infected lungs. However, IAV-induced airspace liquid absorption was overcome to restore, hence “rescue” AWL secretion in IAV-infected lungs by alveolar CFTR activation or potentiation.

Notably, dextran fluorescence was steady in airspaces of CFTR-inhibited alveoli (Figure 3F, right bar) and ENaC-inhibited alveoli (Figure 3G, second bar). These findings indicate CFTR and ENaC inhibition did not reveal, respectively, underlying airspace liquid absorption in untreated lungs, or AWL secretion in IAV-infected lungs. We interpret from these findings that CFTR does not regulate ENaC function in alveolar epithelium, though it may have a regulatory role in other epithelia (39).

### IAV-induced loss of AWL secretion results from alveolar CFTR dephosphorylation

Known mechanisms of CFTR inhibition include CFTR protein degradation (28, 29) and dephosphorylation (40, 41). Immunoblots (Figure 4A) and band densitometry (Figure 4B) show whole lung content of dephosphorylated CFTR protein was higher in IAV- than PBS-instilled lungs, whereas total CFTR protein content was equivalent. These findings indicate IAV induced CFTR dephosphorylation in the lung. Immunofluorescence staining of fixed lung tissue affirms that IAV lung infection induced CFTR dephosphorylation and adds that the dephosphorylation occurred in the alveolar epithelium (Figure 4, C-G).

**Figure 4.**
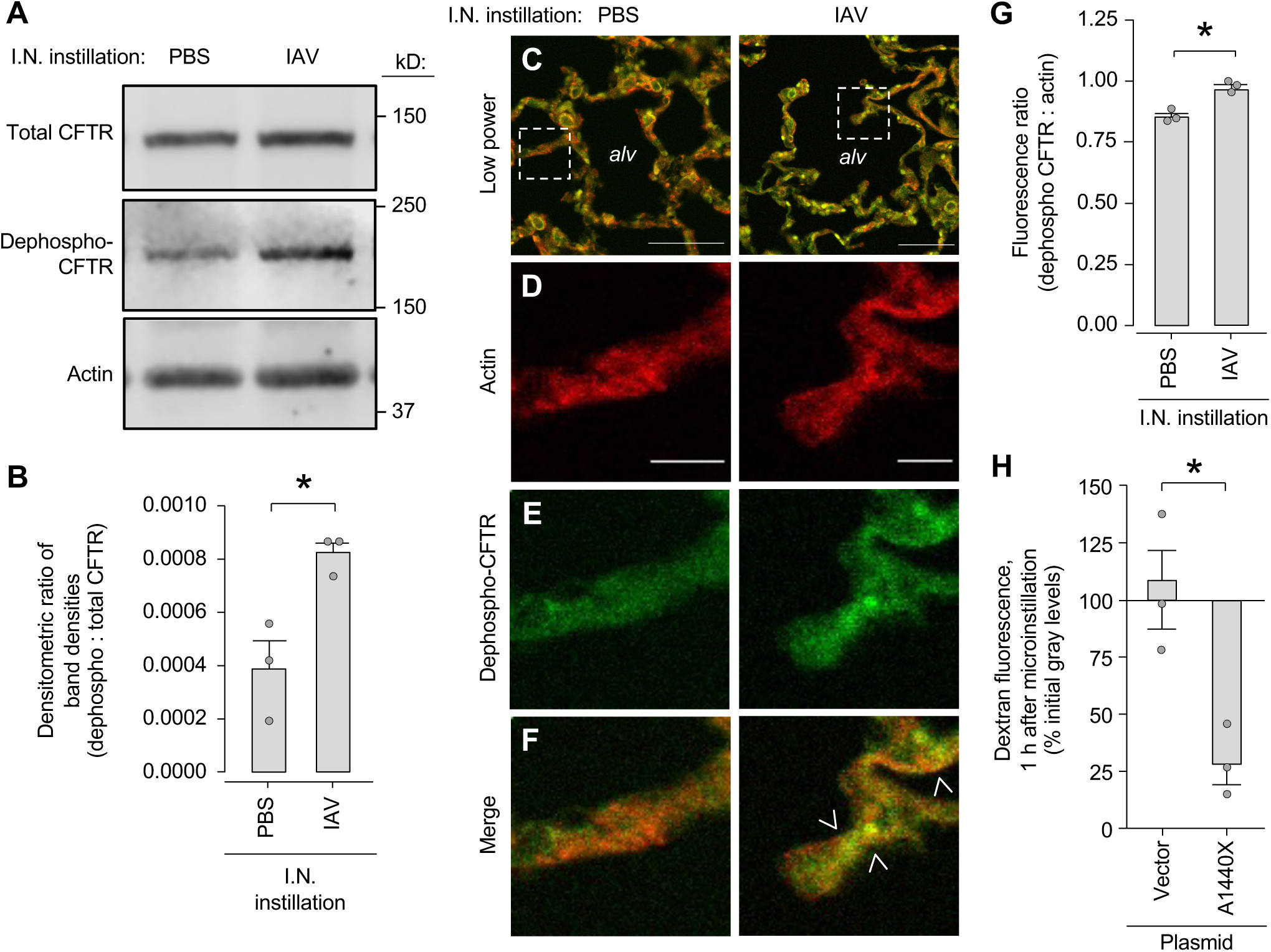
IAV lung infection causes CFTR dephosphorylation to block AWL secretion. Lungs from mice intranasally instilled with IAV or PBS were excised at 24 h post-instillation and homogenized, fixed, or imaged. For group data, circles indicate *n* and each represent one mouse. Bars: mean ± SEM; \**P* < 0.05 by two-tailed *t* test. (**A**-**B**) Representative images (A) and group data (B) show immunoblot results of whole lung homogenate with the indicated antibodies (A). (**C**-**G**) Representative images (C-F) and group data (G) show results of immunofluorescence microscopy using the indicated antibodies on fixed lung tissue. Low (C) and high (D-F) power views show immunofluorescence of representative alveolar septa. *Dashed boxes* (C) indicate septal regions shown in D-F. Note, bright dephospho-CFTR fluorescence (*green*) resulted in yellow pseudocolor (*arrowheads* in F) only in alveoli of IAV-infected lungs (*right column*). For group data (G), mean fluorescences were quantified from images of six randomly selected alveolar fields, each containing at least 30 alveoli. *Alv*, alveolar airspace. Scale bars: 25 (C) and 10 (D) µm. (**H**) Group data derived by confocal imaging of live, intact, perfused mouse lungs show change of TRITC-dextran fluorescence in alveolar airspaces after alveolar dextran microinstillation. Mice were pretreated with intranasal instillation of liposome-complexed plasmid DNA conferring expression of the plasmid vector or A1440X mutant CFTR protein, as indicated, at 24 h prior to IAV instillation and 48 h prior to lung excision for imaging. Mean dextran fluorescence change was quantified in an imaging field of at least 30 alveoli.

To evaluate the role of CFTR dephosphorylation in the mechanism of IAV-induced AWL inhibition, we quantified AWL secretion in lungs pretreated with plasmid DNA conferring expression of dephosphorylation-resistant CFTR protein (40, 42). The A1440X mutant CFTR protein contains a stop mutation at residue 1440, resulting in deletion of the 40 C-terminal amino acids, including major phosphatase binding sites (40, 42). Thus, although the A1440X mutant CFTR retains cell surface expression and Cl^-^ channel activity, it is dephosphorylated at a slow rate (40). Since we have shown previously that intranasal plasmid instillation induces plasmid protein expression in alveolar epithelium (26), we pretreated mice with intranasal instillation of vector or A1440X plasmid. Our findings show airspace dextran fluorescence decreased in alveoli of IAV-infected mice pretreated with the A1440X CFTR plasmid but not vector (Figure 4H), indicating plasmid pretreatment restored AWL secretion in IAV-infected lungs. Taking these findings together with the immunoblot and immunofluorescence data, we interpret IAV lung infection inhibited AWL secretion by inducing CFTR dephosphorylation in alveolar epithelium.

### IAV causes alveolar retention of inhaled SA

Our published data show alveolar epithelial CFTR inhibition blocks spontaneous alveolar clearance of SA (26), raising the possibility that loss of AWL secretion promotes alveolar stabilization of bacteria. To evaluate this possibility, we quantified the effect of AWL inhibition on alveolar stability of GFP-labeled *S. aureus* (strain USA300) in stationary growth phase (SA^GFP^). We selected the stationary growth phase because it may reflect the state of SA at the initiation of SA lung infection, when SA are inhaled from the nasal cavity (43), since bacteria in stationary-like growth phases are susceptible to surface detachment (44). Our findings show alveolar microinstillation of SA^GFP^ into alveoli that were pretreated with either buffer or CFTRinh-172 caused formation of bacterial microaggregates in alveolar niches (Figure 5A, top row). Alveolar washout by buffer microinstillation caused major loss of SA^GFP^ fluorescence in buffer-pretreated alveoli (Figure 5A, left column and Figure 5B, left bar), indicating SA^GFP^ microaggregates were susceptible to alveolar clearance and, therefore, unstable on the alveolar wall. However, washout failed to dislodge SA^GFP^ that were retained in CFTR-inhibited alveoli for 3 h (Figure 5A, right column and Figure 5B, right bar). These findings indicate retention in CFTR-inhibited alveoli caused SA^GFP^ to change phenotype from unstable to stabilized on the alveolar wall. The significance of this phenotypic change is indicated by our published findings, which show stabilized, but not unstable SA induce alveolar damage leading to fatal acute lung injury (26). Taking our new and published findings together, we interpret SA^GFP^ retention in CFTR-inhibited alveoli caused SA^GFP^ to rapidly shift from an unstable, benign phenotype to a stabilized, virulent phenotype capable of inducing alveolar damage.

**Figure 5.**
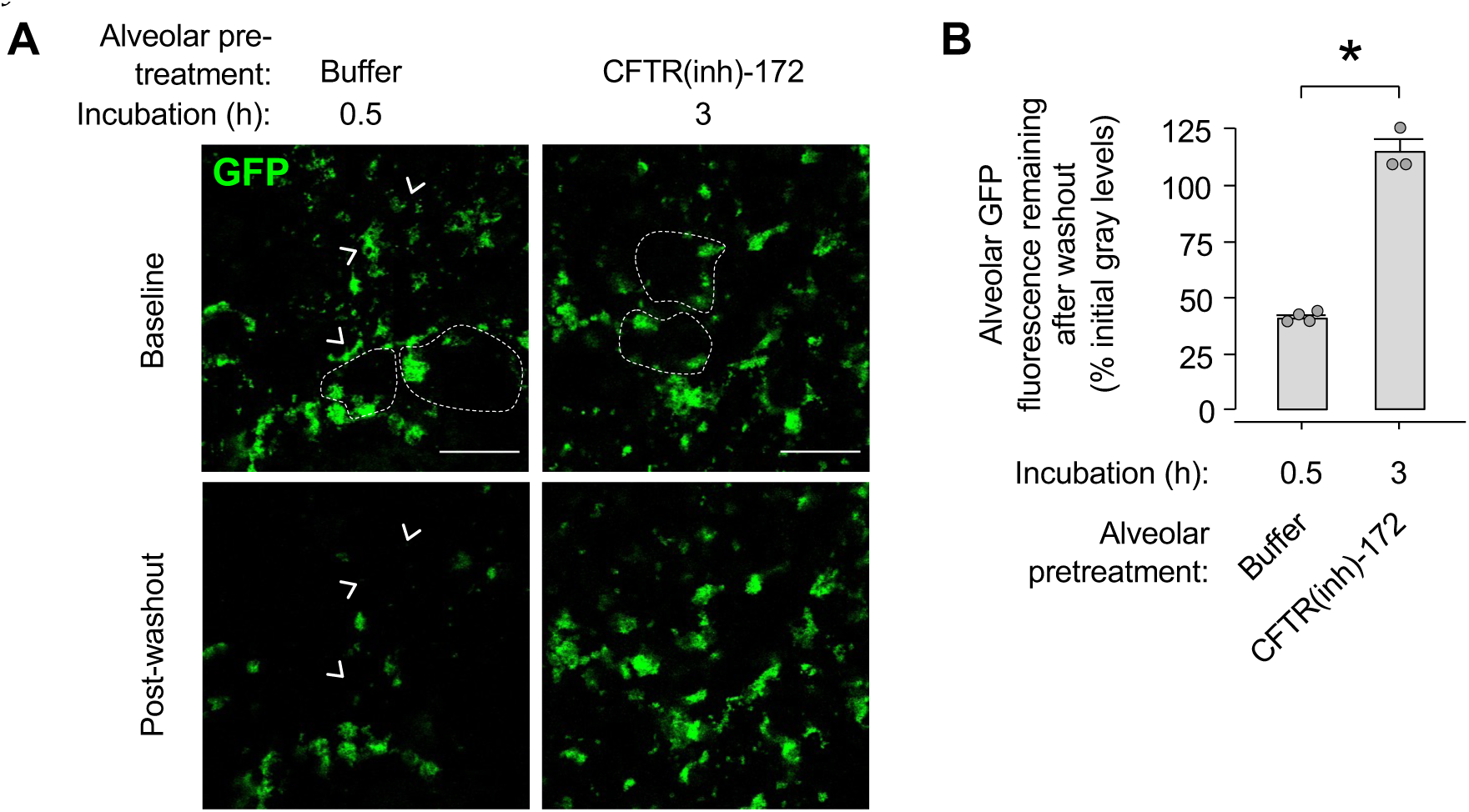
AWL secretion protects against alveolar stabilization of SA^GFP^. Confocal images (**A**) and group data (**B**) show change of SA^GFP^ fluorescence in alveolar airspaces in response to alveolar washout. We pretreated alveoli with microinstillation of CFTR(inh)-172 dissolved in HEPES-buffered solution or buffer alone, as indicated, then we microinstilled SA^GFP^. Alveoli were subjected to washout by alveolar microinstillation of HEPES-buffered solution at 0.5 or 3 h after SA^GFP^ microinstillation as indicated. *Arrowheads* point out example SA^GFP^ microaggregates that had complete loss of fluorescence in response to washout, hence were cleared from alveoli. Fluorescence of the alveolar epithelium is not shown. *Dashed lines* delineate example alveolar walls. Circles (B) indicate *n* and were each generated by comparing mean GFP fluorescence before and after washout in one imaging field of at least 20 alveoli. Bars: mean ± SEM; \**P* < 0.05 by two-tailed *t* test. Scale bars: 50 µm.

Since IAV lung infection blocked alveolar epithelial CFTR function, we considered IAV might promote alveolar retention of inhaled SA^GFP^. To test this hypothesis, we used our live lung imaging approach to view alveoli after two intranasal instillations in mice: IAV or PBS, then 24 h later, SA^GFP^. Within 1 h of instillation in IAV-infected lungs, SA^GFP^ formed groups in alveoli (Figure 6A) that we have previously identified (26). In line with our previous findings (26), small clusters of non-microaggregated SA^GFP^ were present on flat alveolar surfaces (Figure 6A, single arrow), and microaggregates were located at alveolar niches (Figure 6A, inset and double arrows). Microaggregates and small clusters formed with equal frequency (Figure 6B) and size (Figure 6C) in alveoli of mice pretreated with IAV or PBS, indicating the micromechanical features of alveoli that determine bacterial group formation were preserved in IAV-infected lungs. To determine the time course of spontaneous bacterial clearance, we imaged SA^GFP^-containing alveoli at 1 and 3 h after SA^GFP^ instillation. Whereas most SA^GFP^ groups in PBS-pretreated lungs had complete loss of alveolar fluorescence (Figure 6D, left column, arrowheads and 6E, left bar), indicating the bacteria were spontaneously removed from alveoli, the rate of alveolar SA^GFP^ fluorescence loss was markedly diminished in IAV-infected lungs (Figure 6D, right column and 6E, right bar), indicating IAV caused failure of alveolar SA^GFP^ clearance. Taking these findings together, we interpret that while IAV lung infection had no effect on niche-based microaggregation by inhaled SA^GFP^, the infection blocked alveolar SA^GFP^ clearance, causing retention of inhaled SA^GFP^ in alveoli.

**Figure 6.**
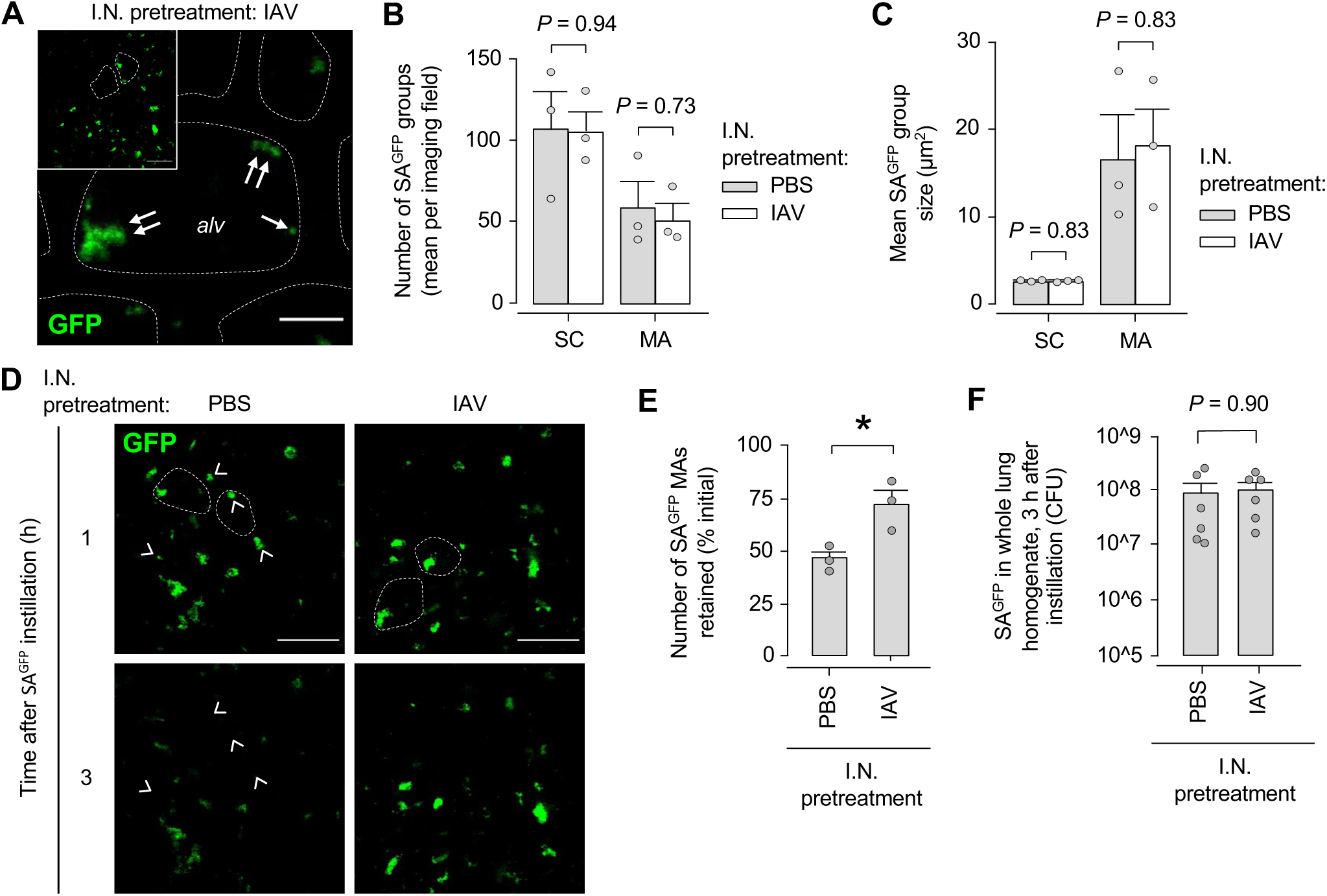
IAV lung infection causes alveolar retention of inhaled SA^GFP^. Mice were pretreated with intranasal instillation of IAV or PBS as indicated, then intranasally instilled with SA^GFP^ 24 h later. For group data, circles indicate *n* and each represent one mouse. Bars: mean ± SEM; *P* calculated by two-tailed *t* test. (**A-C**) Low (*inset*) and high power confocal images (A) show SA^GFP^ fluorescence in live alveoli of intact, blood-perfused, IAV-infected mouse lungs, 1 h after intranasal SA^GFP^ instillation. *Dashed lines* delineate example alveolar walls (fluorescence not shown). *Single* and *double arrows* indicate, respectively, SA^GFP^ grouped as small clusters (*SC*) and microaggregates (*MA*). Group data show number (B) and size (C) of SCs and MAs in alveoli of lungs pretreated with PBS or IAV instillation as indicated. For B and C, SA^GFP^ group number and size were quantified as means in at least two imaged fields of 30 alveoli each. *Alv*, example alveolar airspace. Scale bars: 50 (*inset*) and 10 µm. (**D-E**) Confocal images (D) show alveolar SA^GFP^ fluorescence at 1 h (*top row*) and, in the same alveoli, at 3 h (*bottom row*) after SA^GFP^ instillation. *Arrowheads* indicate example MAs that spontaneously lost all fluorescence, hence were cleared from alveoli. Group data (E) show the proportion of SA^GFP^ MAs that maintained alveolar fluorescence, hence were retained in alveoli. For E, MAs were quantified as the mean proportion retained in at least two imaged fields of 30 alveoli each. Scale bars: 50 µm. (**F**) Content of viable SA^GFP^ in lung homogenate at 3 h after intranasal SA^GFP^ instillation.

IAV-induced inhibition of AWL secretion might cause alveolar SA^GFP^ retention by loss of convective particle transport or impairment of bacterial killing. To evaluate the possibility that the retention resulted from bacterial killing impairment, we quantified lung content of viable SA^GFP^. Recovery of an equal number of viable SA^GFP^ from whole lung homogenate of mice pretreated with IAV and PBS (Figure 6F) indicates bulk mechanisms of bacterial killing were intact in IAV-infected lungs. Since neutrophils are major effectors of bacterial killing in the lung (45), we quantified neutrophils in alveolar airspaces by alveolar microinjection of fluorophore-labeled antibodies against CD11b (46), a surface protein expressed by neutrophils and other innate immune cells in the lung (47). CD11b fluorescence was absent in airspaces at 3 h after intranasal PBS instillation, 24 h after IAV instillation, and 3 h after SA^GFP^ instillation, but it increased markedly at 9 h after SA^GFP^ instillation (Supplemental Figure 2, A-E). These findings show that although intranasal SA^GFP^ instillation stimulated airspace recruitment of CD11b+ cells, the recruitment was delayed for hours. Since CD11b+ cells were absent from airspaces during the time we observed IAV-induced alveolar SA^GFP^ retention, we rule out a role for CD11b+ cells in the retention mechanism. Together, these findings suggest bacterial killing impairment did not explain IAV-induced alveolar SA^GFP^ retention and support the possibility that the retention resulted from failure of convective bacterial transport.

### AWL rescue protects against Hla-induced alveolar damage

We have reported SA-induced acute lung injury results from bacterial secretion of the membrane pore-forming toxin, alpha hemolysin (Hla), leading to alveolar epithelial membrane damage, alveolar barrier loss, and fatal lung injury (26). We considered alveolar retention of SA^GFP^ in IAV-infected lungs might augment Hla-induced alveolar damage (Figure 7, A-D). To test this hypothesis, we evaluated epithelial membrane damage in terms of epithelial calcein fluorescence, which decreases following membrane damage (26, 48). Since we have shown the extent of Hla-induced alveolar epithelial damage is Hla dose-dependent (26), we selected an Hla dose that failed to induce epithelial calcein fluorescence loss after microinstillation into alveoli of unchallenged lungs (Figure 7, A and D, left bar). However, the microinstillation induced major calcein loss in alveoli of IAV-infected lungs (Figure 7, B and D, middle bar), indicating IAV lung infection augmented Hla’s membrane-damaging effect. Because alveolar pretreatment with ivacaftor blocked the calcein loss (Figure 7, C and D, right bar), we suggest IAV-induced CFTR dysfunction, hence AWL inhibition underpinned the augmented alveolar damage response. These findings support the notion that IAV-induced inhibition of AWL secretion increased alveolar epithelial damage by secreted SA toxin, perhaps by impairing toxin clearance from alveoli.

**Figure 7.**
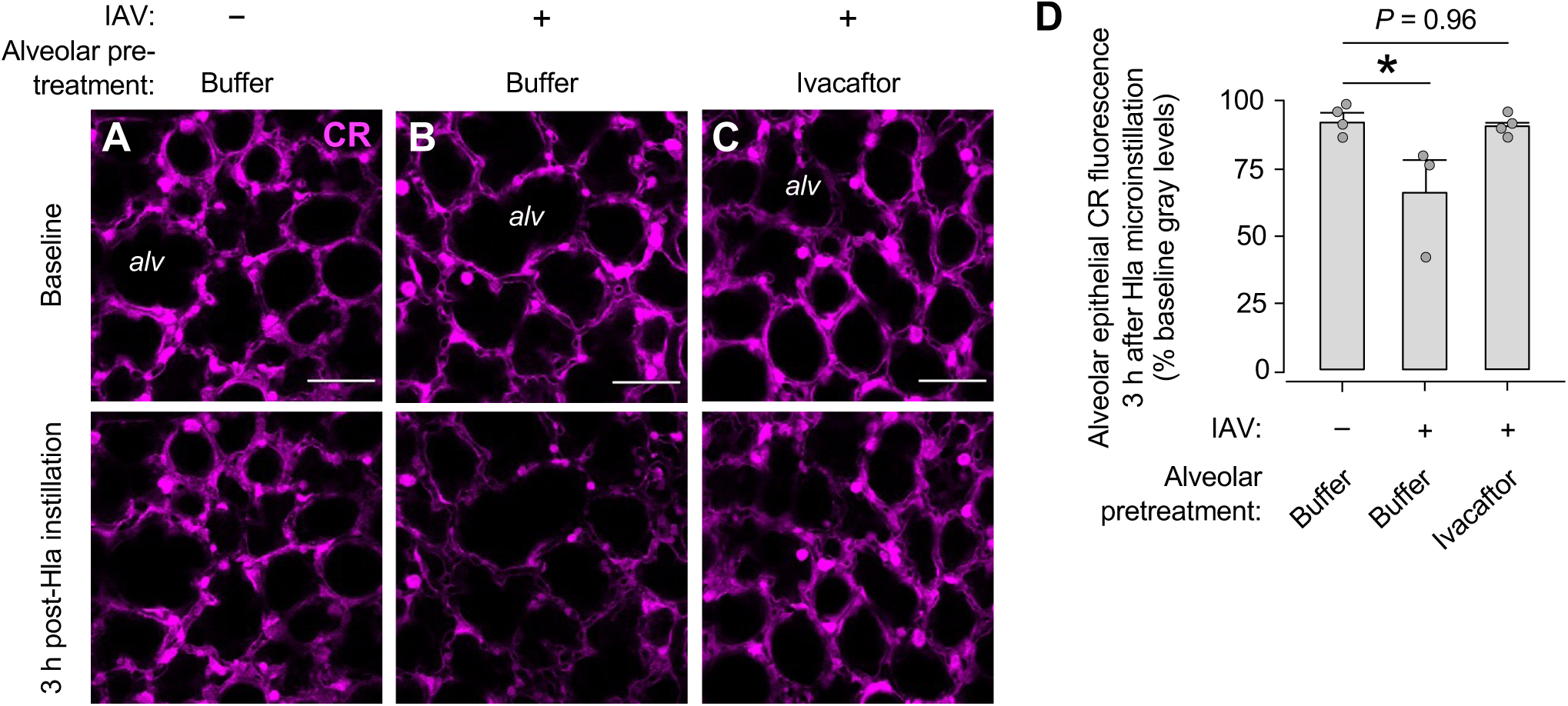
AWL rescue protects against Hla-induced alveolar damage. Confocal images (**A-C**) and group data (**D**) show change of alveolar epithelial calcein fluorescence in response to alveolar microinstillation of purified Hla toxin. Imaged lungs were untreated (-) or intranasally instilled with IAV at 24 h prior to imaging (+) as indicated. Alveolar epithelium was pretreated with alveolar microinfusions of calcein red-orange (*CR*), then ivacaftor dissolved in HEPES-buffered solution or buffer alone, as indicated, before Hla microinstillation. Circles (D) indicate *n* and were each generated by quantifying mean CR fluorescence in one imaging field of at least 30 alveoli at 5 min before and 3 h after Hla microinstillation. Bars: mean ± SEM; \**P* < 0.05 by ANOVA with post hoc Tukey testing. *Alv,* alveolar airspace. Scale bars: 50 µm.

### Efficacy of AWL rescue therapy in mouse models

After confirming systemic ivacaftor injection rescued AWL secretion in IAV-infected lungs (Figure 8A), we tested the therapeutic potential of AWL rescue against SA^GFP^-induced acute lung injury in a mouse model of IAV-SA lung coinfection (Figure 8B). Whereas mice infected with either IAV or SA^GFP^ survived at least 4 days after IAV or SA^GFP^ instillation (data not shown), coinfected mice treated with vehicle injections had high mortality (Figure 8C, solid line), in line with reports by others (21, 49). By contrast, all coinfected mice treated with ivacaftor survived (Figure 8C, dashed line). We interpret IAV augmented SA^GFP^ pathogenesis to induce mortality, but the mortality was decreased by systemic CFTR potentiation therapy.

**Figure 8.**
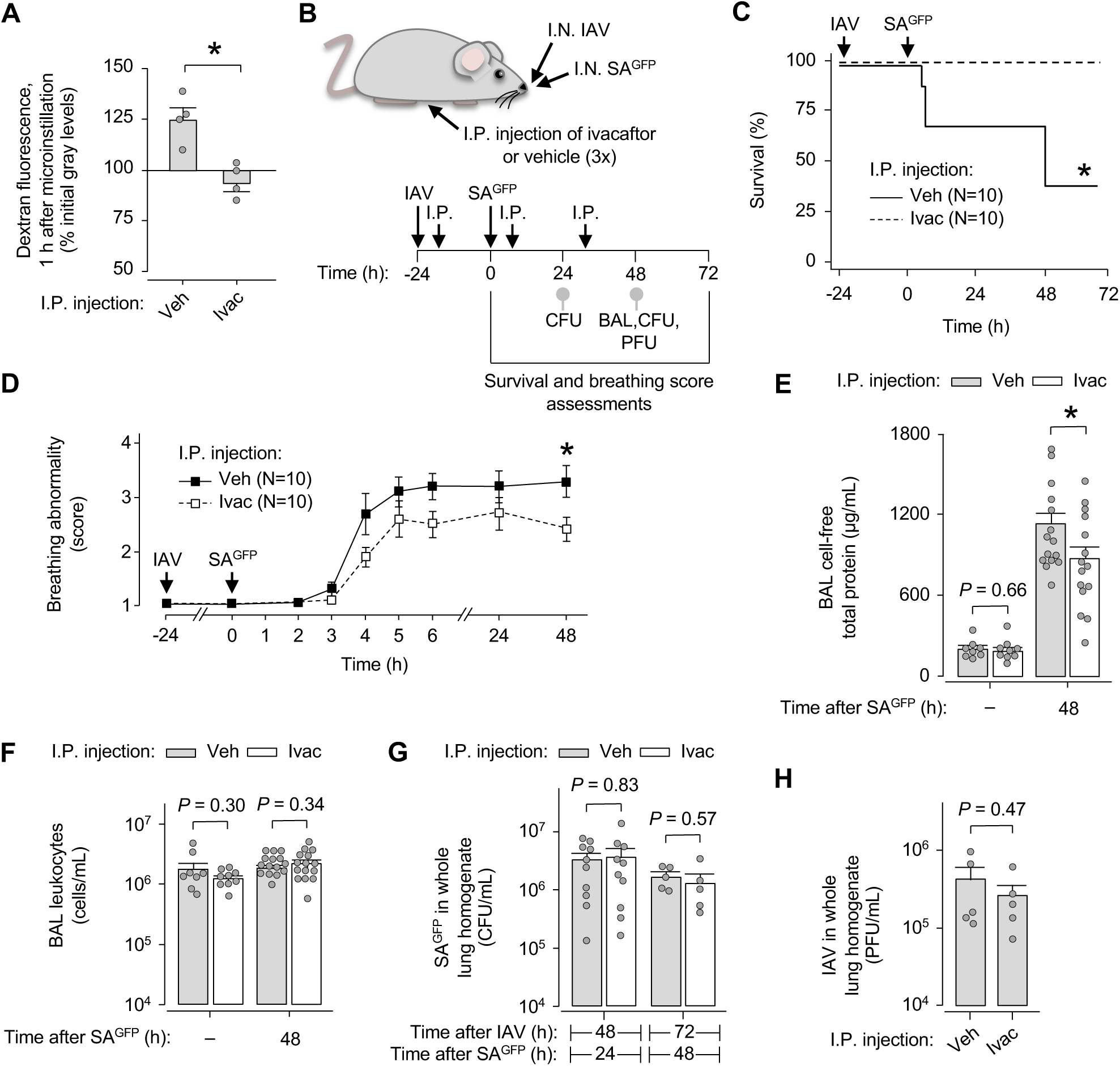
AWL rescue therapy protects against fatal IAV-SA^GFP^ coinfection. (**A**) Group data show change of TRITC-dextran fluorescence in alveolar airspaces after alveolar dextran microinstillation. Mice were given intranasal instillation of IAV, then at 6 h, intraperitoneal (*I.P.*) injection of vehicle (*veh*) or ivacaftor (*ivac*) as indicated. Lungs were excised for imaging at 24 h after IAV instillation. Circles indicate *n* and each represent one mouse in which mean dextran fluorescence change was quantified in an imaging field of at least 30 alveoli. Bars: mean ± SEM; \**P* < 0.05 by two-tailed *t* test. (**B-H**) Experimental design (B) for group data shown in C-H shows timing of intranasal (*I.N.*) instillations, intraperitoneal injections, and procedures including mouse survival (C) and breathing score (D) assessments, bronchoalveolar lavage (*BAL*) fluid collection for quantification of total protein (E) and leukocyte (F) content, and lung excision for quantification of SA^GFP^ (*CFU*; G) and IAV (*PFU*; H) content. Some mice were intranasally instilled with IAV but not SA^GFP^ ((-) in E and F, and all data in H). Breathing scores (D) were imputed for non-surviving mice using their last observed value. Squares (D) and bars (F-H) indicate mean ± SEM; circles (E-H) indicate *n* and each represent data from one mouse; \**P* < 0.05 by log rank (C) or two-tailed *t* test (D-H) as indicated. BAL content of protein (E) and leukocytes (F) were quantified using the same BAL fluid specimen.

Systemic CFTR potentiation might promote survival of coinfected mice by AWL rescue, hence inhibition of alveolar SA^GFP^ retention and SA^GFP^-induced alveolar damage, or alternative mechanisms such as drug-induced bacterial killing (50) or alteration of leukocyte proteins (51) that bear on lung immune recruitment. We addressed these possibilities by quantifying the extent to which systemic CFTR potentiation determined pulmonary edema responses, immune responses, and pathogen burden after IAV and SA^GFP^ instillation (Figure 8B). To determine responses to SA^GFP^, we first assigned scores to IAV- and SA^GFP^-instilled mice using an observational breathing score system (Supplemental Figure 3A) in which higher score correlated with higher bronchoalveolar lavage (BAL) fluid protein content (Supplemental Figure 3B), a marker of alveolar fluid barrier dysfunction, thus pulmonary edema responses. Within hours of instillation, SA^GFP^ induced breathing abnormalities that persisted for days and were decreased in ivacaftor-treated mice (Figure 8D). In line with the breathing scores, direct measures of BAL protein content show SA^GFP^ caused major increase of BAL protein that was markedly reduced by ivacaftor (Figure 8E), indicating ivacaftor protected against SA^GFP^-induced pulmonary edema. By contrast, ivacaftor failed to affect BAL leukocyte content (Figure 8F) or lung content of viable SA^GFP^ (Figure 8G) in coinfected mice and had no impact on lung content of IAV in mice instilled with IAV alone (Figure 8H). These findings indicate systemic CFTR potentiation protected against SA^GFP^-induced pulmonary edema responses in IAV-infected mice without altering lung pathogen burden or number of airspace leukocytes accessible by BAL. Taking these findings together with the survival data, we interpret the survival benefit of systemic CFTR potentiation in coinfected mice as having resulted from protection against SA^GFP^-induced alveolar barrier dysfunction, and we suggest the protection stemmed from alveolar effects of systemic CFTR potentiation therapy, namely AWL rescue.

In a proof-of-concept experiment, we tested the therapeutic potential of AWL rescue by intranasally instilling mice with vector or A1440X plasmid, which restored AWL secretion in IAV-infected mice (Figure 4H), then IAV and SA^GFP^ (Figure 9A). Whereas vector-treated mice had high mortality (Figure 9B, solid line), the mortality (Figure 9B, dashed line) and breathing scores (Figure 9C, dashed line) were markedly decreased in mice treated with A1440X. These findings indicate lung epithelial expression (26) of dephosphorylation-resistant CFTR protein protected against SA^GFP^-induced mortality in IAV-infected mice, supporting the notion that IAV-induced CFTR dephosphorylation in alveolar epithelium was central to the lung pathogenesis of IAV-SA^GFP^ coinfection.

**Figure 9.**
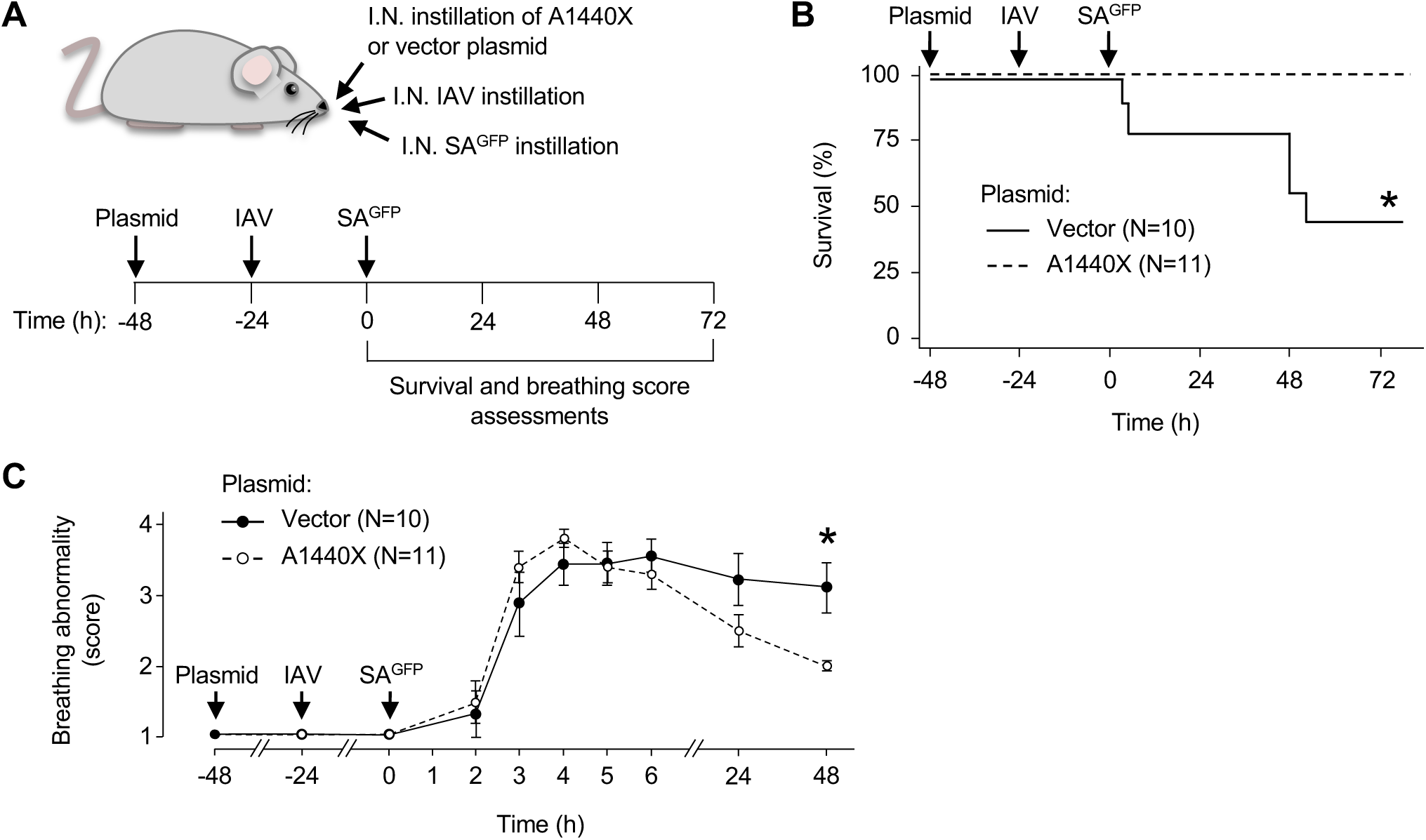
Protection against SA^GFP^-induced mortality in IAV-infected mice by mutant CFTR protein. Experimental design (**A**) for group data shown in **B** and **C** indicates timing of intranasal instillations and survival (B) and breathing score (C) assessments. Breathing scores were imputed for non-surviving mice using their last observed value. Circles (C) indicate mean ± SEM; *n* as indicated; \**P* < 0.05 by log rank testing (B) or two-tailed *t* test (C).

## DISCUSSION

Our findings show, for the first time to our knowledge, that IAV lung infection caused reversal of alveolar liquid dynamics from epithelial secretion to absorption. The reversal resulted from two effects of IAV lung infection on alveolar epithelium: dephosphorylation-mediated CFTR inhibition, leading to loss of AWL secretion; and ENaC stimulation, inducing alveolar liquid absorption. Alveolar epithelial treatment with CFTR activator or potentiator drugs rapidly restored AWL secretion in IAV-infected lungs, indicating the electrochemical and osmotic gradients that normally drive AWL secretion were intact, and thus the AWL inhibition resulted directly from CFTR inhibition. The effect was to abrogate a major mechanism of alveolar defense, namely AWL secretion, to generate an alveolar microenvironment hospitable to SA stabilization and Hla-induced epithelial damage. Thus, the new understanding gained from our findings is that AWL secretion is a major mechanism of innate alveolar defense against inhaled bacteria. Its inhibition by IAV lung infection promoted initiation of SA^GFP^ infection and SA^GFP^-induced tissue damage in alveoli, leading to fatal SA^GFP^-induced lung injury.

Since acute lung injury is defined by *gain* of airspace liquid, as pulmonary edema fluid (12, 13), our proposal that coinfection-induced lung injury resulted from *loss* of airspace liquid, as AWL, adds new understanding to lung injury mechanisms. Our findings show IAV lung infection inhibited AWL secretion, leading to retention of SA^GFP^ against the alveolar wall for hours. The retention enabled SA^GFP^, a pathogen known for its ability to rapidly adapt to its microenvironment (52), to transition from an unstable phenotype susceptible to alveolar clearance to a highly stable phenotype that resisted alveolar dislodgement. Thus, the alveolar microenvironment generated by IAV-induced loss of AWL secretion created an opportunity for SA^GFP^ to rapidly assume a stabilized phenotype that, we have shown previously (26), initiates alveolar infection and fatal alveolar damage. Thus, we interpret AWL secretion contributes critically to alveolar defense against inhaled bacteria, and AWL inhibition constitutes a major mechanism by which IAV promotes secondary bacterial infection.

How IAV-induced AWL inhibition caused alveolar SA^GFP^ retention remains unclear. Since AWL secretion induces flow of liquid along the alveolar wall that convectively transports particles out of alveoli (31), AWL inhibition might block convective, alveolus-to-airway transport that normally clears alveoli of bacteria. Alternatively, AWL inhibition might impair bacterial killing by altering alveolar surfactant activity, macrophage function, or airspace liquid pH. Since our findings show IAV lung infection blocked loss of SA^GFP^ fluorescence in alveoli but did not alter SA^GFP^ counts in the whole lung, and we did not identify neutrophils in alveolar airspaces during the period of observed alveolar SA^GFP^ retention, failure of bacterial killing probably did not underlie alveolar SA^GFP^ retention. Rather, we suggest the retention was caused by loss of convective bacterial transport.

Our findings show IAV lung infection caused CFTR dephosphorylation in alveolar epithelium. It is known that CFTR is expressed by alveolar epithelial type 1 and 2 cells (53–58) and drives alveolar epithelial secretion of Cl^-^ and liquid under baseline conditions (31, 56, 59–61). Here, we add that IAV lung infection blocked CFTR-mediated AWL secretion and induced alveolar epithelial CFTR dephosphorylation. Plasmid-mediated inhibition of dephosphorylation restored AWL secretion in IAV-infected lungs, indicating epithelial CFTR dephosphorylation was responsible for AWL inhibition. The causative role of CFTR dephosphorylation in IAV-induced AWL inhibition is further supported by the efficacy of ivacaftor for restoring AWL secretion, since ivacaftor’s capacity to potentiate function of wild type CFTR is increased in cells exposed to non-phosphorylating conditions (62). Thus, our findings reveal a new mechanism of IAV-induced CFTR inhibition in the lung. Although others have shown IAV induces CFTR ubiquitination and degradation in cultured airway epithelial cells (28–30), we do not identify that IAV-induced CFTR dephosphorylation has been reported previously. The extent to which IAV induces CFTR dephosphorylation versus other CFTR inhibition mechanisms, and whether the mechanisms differ among lung epithelia, remain unclear. Factors that might determine epithelium-specific CFTR inhibition mechanisms include cellular patterns of CFTR expression (53, 58) and processing (63, 64), the degree to which epithelial cells are directly infected (65), and the extent to which uninfected epithelium communicates with infected cells (26, 46, 66, 67). These issues, as well as the upstream mechanisms responsible for IAV- induced CFTR dephosphorylation, warrant further study.

Our finding, that IAV lung infection induced a switch of alveolar liquid dynamics from basal AWL secretion to airspace liquid absorption, is novel but supported by existing literature. Our data demonstrating basal AWL secretion aligns with reports indicating the intact alveolar epithelium secretes Na^+^ (59), Cl^-^ (31, 59, 60), and liquid (31, 59) under baseline conditions. Although it is known that alveolar epithelial type 1 and 2 cells express ENaC (35, 36, 68) and exhibit rapid reversal from liquid secretion to absorption in response to local conditions (56, 59), our findings are the first, to our knowledge, to show that reversal from CFTR-mediated liquid secretion to ENaC-mediated liquid absorption occurred in alveolar epithelium in response to IAV lung infection. The reversal had major bearing on the outcome of IAV-SA coinfection infection in that it promoted alveolar SA^GFP^ retention and Hla-induced alveolar epithelial damage. Notably, others have reported seemingly opposite findings, that IAV blocks lung absorption of airway-instilled liquid in rodents (67, 69). However, we point out that the contribution of the small airway epithelium to bulk transport of airway-instilled liquid is unclear and may be a determinant of IAV-induced impairment of liquid absorption after airway instillation. Our findings using the live lung imaging approach provide direct evidence that IAV lung infection induced airspace liquid absorption in intact, perfused alveoli.

From a therapeutic perspective, our data suggest AWL rescue represents a new therapeutic target for preventing fatal SA-induced acute lung injury after IAV infection. Our findings show AWL rescue by two approaches – systemic administration of the CFTR potentiator drug, ivacaftor, and intranasal instillation of plasmid DNA conferring dephosphorylation-resistant CFTR protein expression – each protected against SA^GFP^-induced mortality in IAV-infected mice. Importantly, ivacaftor decreased SA^GFP^-induced alveolar barrier dysfunction without altering BAL-accessible lung leukocyte numbers or lung pathogen burden, placing alveolar responses to IAV and SA^GFP^ at the center of coinfection pathogenesis mechanisms and suggesting improvement of alveolar function accounted for ivacaftor’s therapeutic effect. Taking these findings together with the imaging data, we propose fatal SA^GFP^-induced acute lung injury resulted from IAV-induced loss of AWL secretion, leading to alveolar retention of SA^GFP^ and augmenting Hla-induced alveolar damage. Approaches that rescue AWL secretion in IAV-infected lungs, perhaps by blocking alveolar epithelial CFTR dephosphorylation, may prevent secondary SA^GFP^ infection in alveoli, thereby decreasing SA^GFP^-induced lung injury and mortality. Since ivacaftor is already in clinical use and has an excellent safety and tolerability profile (70), these findings may be translatable to patients.

In conclusion, our findings show IAV lung infection induced reversal of normal alveolar liquid dynamics to cause airspace liquid absorption and inhibit AWL secretion. Thus, IAV infection abrogated a major mechanism of alveolar defense, leading to alveolar retention of inhaled SA^GFP^, Hla-induced alveolar damage, and fatal SA^GFP^-induced acute lung injury. Rescue of AWL secretion was protective. These findings contribute new understanding of the role of alveolar liquid in health and lung injury. First, AWL secretion was critical to lung defense against inhaled bacteria, since loss of AWL secretion promoted alveolar SA^GFP^ infection. Second, although pathogen-induced lung injury is traditionally defined by gain of alveolar liquid (edema), our findings show loss of alveolar liquid (AWL) was critical to IAV lung pathogenesis. We propose therapeutic approaches that restore AWL secretion in IAV-infected lungs may protect against fatal secondary SA lung infection.

## METHODS

### Experimental design

Experiments were designed in line with ARRIVE guidelines. Experimental units were single mice unless otherwise indicated in legends. Mice used for survival assessments, BAL fluid collection, or lung excision for pathogen quantification were allocated to groups in a manner that ensured roughly equal mean mouse weight per group. Groups were mixed within cages, and assessments of breathing abnormalities and need for euthanasia were carried out by an investigator blinded to mouse groups. In rare instances, mice were excluded from analyses if instillation of pathogens, plasmid, or drugs failed to meet our laboratory’s standards for quality. Exclusion decisions were made in an identical manner for all groups and based on instillation quality assessments that were completed at the time of the instillations, thus prior to experiment completion and analysis. Outcome measures are indicated in figures and legends.

### Fluorophores

We purchased calcein AM (10 μM), calcein red-orange AM (10 μM), and tetramethylrhodamine (TRITC)-conjugated dextran (10 mg/mL) from ThermoFisher.

### Reagents

Forskolin (20 uM) and amiloride (10 uM) were purchased from Selleck Chemicals and CFTR(inh)-172 (20 uM) from Sigma. Reagents were freshly constituted for experiments. Ivacaftor was purchased from Selleck Chemicals and reconstituted on delivery with DMSO (Fisher Scientific) to a concentration of 50 mg/mL before aliquoting and storing at -80C. We prepared single doses of ivacaftor or vehicle within 1 h of administration. Doses for intraperitoneal injection were 40 mg/kg ivacaftor in 5% DMSO (ThermoFisher), 5% Tween 80 (Selleck Chemicals), 40% PEG300 (Selleck Chemicals), and 50% of 0.9% saline (Grifols) solution. Vehicle was the identical weight-based solution volume without ivacaftor. Purified Hla (1 mg/mL at 2332 HU/mg, diluted 1:10 in HEPES-buffered solution as below) was a gift of Dr. Juliane Bubeck-Wardenburg’s laboratory (Washington University School of Medicine).

### Solutions

We purchased Ca^2+^- and Mg^2+^-containing DPBS and Ca^2+^- and Mg^2+^-free PBS from Corning. Isolated lungs were perfused with HEPES-buffered vehicle solution of pH 7.4 and osmolarity 295 mOsm and containing 150 mM Na^+^, 5 mM K^+^, 1 mM Ca^2+^, 1 mM Mg^2+^, and 10 mM glucose. Except where noted, fluorophores, reagents, and antibodies microinstilled in alveoli were dissolved or suspended in the same HEPES-buffered solution.

### Antibodies

Antibodies were purchased from commercial vendors or academic institutions that provided antibody validation information. Allophyocyanin (APC)-conjugated monoclonal Ab against CD11b (clone M1/70; ThermoFisher) was diluted 1:25 for alveolar microinstillation. Antibodies used for immunoblotting and immunofluorescence microscopy include: mouse monoclonal Ab against CFTR (clone A-3; Santa Cruz Biotechnology); mouse monoclonal Ab against dephosphorylated CFTR (clone 570; University of North Carolina at Chapel Hill CFTR Antibody Distribution Program); and rabbit polyclonal Ab against actin (Sigma). For immunoblotting, antibodies were diluted in StartingBlock T20 (TBS) Blocking Buffer (ThermoFisher) and incubated with membranes as follows: CFTR mAb A-3 was diluted to 1:100 and incubated for 24 h at 4°C; CFTR mAb 570 was diluted to 1:500 and incubated for 24 h at 4°C; and actin Ab was diluted to 1:2,000 and incubated for 1 h at room temperature. For immunofluorescence microscopy, CFTR mAb 570 was diluted to 1:250 in 0.2% Triton-X-100 (ThermoFisher) and 2% Bovine Serum Albumin (ThermoFisher) and incubated with specimens for 24 h at 4°C, and actin Ab was diluted to 1:50 in 0.2% Triton-X-100 and incubated for 24 h at 4°C. Fluorophore-tagged secondary antibodies included goat anti-mouse IgG (Alexa Fluor 488) and goat anti-rabbit IgG (Alexa Fluor 647) and were purchased from Jackson, diluted to 1:50 in 0.2% Triton-X-100, and incubated with specimens for 1 h at room temperature.

### Viral strain, preparation, and inoculation

Mouse-adapted IAV A/Puerto Rico/8/934 (H1N1) was propagated in 8-day old embryonated chicken eggs (Charles River Laboratories), diluted in DPBS containing Ca^2+^ and Mg^2+^, aliquoted immediately after propagation, and stored at -80°C until thawing for instillations. IAV was intranasally instilled in anesthetized mice within 1 h of thawing at an inoculation dose of 2000 PFU in 20 uL or, for selected imaging experiments, 5000 PFU in 50 uL.

### Bacterial strain, preparation, and inoculation

We used GFP-tagged *S. aureus* strain USA300 LAC (SA^GFP^) for all experiments involving bacteria. Bacteria were stored at -80°C in 25% glycerol in autoclaved Luria-Bertani (LB) broth media (MP Biomedicals) and propagated on LB-agar plates containing chloramphenicol (10 ug/mL). Plates were refreshed from frozen stock every 1-2 weeks. For experiments, single bacterial colonies were propagated in autoclaved LB media containing chloramphenicol (10 ug/mL) in a shaking incubator at 37°C and 250 rpm (New Brunswick Scientific) for 18 h. Bacteria were prepared for alveolar microinstillation or intranasal instillation, respectively, by diluting 1 mL of culture in 500 uL or 1.3 mL in 300 uL of DPBS containing Ca^2+^ and Mg^2+^. Within 40 min of bacterial removal from the incubator, we instilled the bacteria-containing solution into lung alveolar airspaces by alveolar micropuncture (details below) or into mice by intranasal instillation (30 uL to deliver 1 x 10^8^ CFU per mouse). For intranasal instillations, mice were rapidly anesthetized and instilled in pairs to ensure similarity of inocula across animals.

### Animals

Mice were male Swiss Webster, purchased from Charles River Laboratories and Taconic Biosciences, 27-38 g, and 4-8 weeks old. We anesthetized mice using inhaled isoflurane (4%) and intraperitoneal injections of ketamine (up to 100 mg/kg) and xylazine (up to 5 mg/kg) for intranasal instillations and surgical procedures. Mice were anesthetized with isoflurane for intraperitoneal injections. For surgical procedures, we injected the tail veins of anesthetized mice with heparin (50 units; Mylan), then exsanguinated the mice by cardiac puncture.

### Intranasal instillation and intraperitoneal injection

Instillation and injection qualities were recorded on a 4-point scale at the time of instillation or injection by the investigator performing the procedure. In general, quality was considered acceptable for experiments if the instillation was recorded as 3-4 (i.e., little or no loss of instillate observed) or the injection was recorded as 4 (i.e., no injury or injectate fluid leakage at the injection site).

### Isolated, blood-perfused lungs

Using our reported methods (26), we cannulated the trachea, pulmonary artery, and left atrium of the heart of exsanguinated mice, then excised the heart, lungs, and cannulas en bloc. Lungs were inflated with room air through the tracheal cannula and perfused through the pulmonary arterial and left atrial cannulas at 0.4-0.6 mL/min with autologous blood diluted in a solution of 4% dextran (70 kDa; Molecular Probes), 1% FBS (Gemini Bio-Products), and HEPES-buffered solution at pH 7.4, osmolality 320 mOsm/kg, and 37°C. We used in-line pressure transducers (ADInstruments) to maintain constant airway pressure 6 cm H_2_O via a continuous positive airway pressure machine (Philips Respironics) and pulmonary artery and left atrial pressures 10 and 3 cm H_2_O, respectively, via a roller pump (Ismatec). In general, the lungs were positioned to enable micropuncture and imaging of the diaphragmatic surface of the right caudal lobe. Portions of the lung surface that were not used for micropuncture and imaging were covered with plastic wrap to prevent desiccation.

### Alveolar microinstillation

We generated hand-beveled glass micropipettes (Sutter Instruments) in our laboratory to micropuncture single alveoli under bright-field microscopy, as we have done previously (26). Micropunctured alveoli were instilled with fluorophores, reagents, and antibodies in solution, resulting in their spread from the micropunctured alveolus to neighboring alveoli. Additional detail is included in the main text. For bacterial microinstillations, we prepared SA^GFP^-containing solutions as above, then microinstilled the solutions to deliver approximately 10^4^ CFU in three seconds of discontinuous microinstillation (71). Microinstillations were performed in 1-3 alveolar sites bordering each imaging field.

### Live lung imaging and analysis

By our established methods (26), we viewed alveoli by confocal microscopy (LSM800; Zeiss) with a 20x water immersion objective (NA 1.0; Zeiss) and coverslip. We used bright-field microscopy to randomly select regions of 30-50 alveoli for microinstillation and imaging. All images were acquired as single images using Zen (v.2.6; Zeiss) and recorded as Z-sections. Analyzed images were 4-8 um below the pleura. Optical thickness was 32-34 um, and frame size was 512 x 512 pixels. We established laser, filter, pinhole, and detector settings at the beginning of each imaging experiment to optimize alveolar fluorescence and avoid fluorescence saturation, then maintained the settings for the duration of the experiment. We confirmed absence of bleed-through between fluorescence emission channels in experiments using fluorophores of different excitation spectra. Images were analyzed using ImageJ (NIH; v.2.0.0-rc-69/1.52n). Brightness and contrast adjustments were applied to individual color channels of entire images and equally to all experiment groups. We did not apply further downstream processing or averaging.

### Airspace dextran fluorescence determination

We used an established approach (26, 31) to microinstill airspaces with calcein-AM (10 uM), then TRITC-conjugated dextran (70 kDa; 10 mg/mL in HEPES-buffered solution). Provided alveolar fluid barrier function is intact, time-dependent loss of TRITC-dextran fluorescence indicates dilution by AWL secretion (31).

### Alveolar permeability determination

To determine alveolar barrier properties, we added TRITC-conjugated dextran (70 kDa; 10 mg/mL) to the intact lung perfusate solution using our established methods (26). Normal permeability of the alveolar fluid barrier was detected by absence of TRITC-dextran fluorescence in alveolar airspaces (26).

### Immunoblot

Using our reported methods (26), we cannulated the trachea and pulmonary artery of exsanguinated mice, then washed the lung vessels with ice-cold DPBS containing Ca^2+^ and Mg^2+^. The lungs were excised and snap-frozen in liquid nitrogen, then pulverized inside a specimen bag using a mortar and pestle. The pulverized lungs were mixed and incubated with RIPA buffer (ThermoFisher) and Halt protease mix (ThermoFisher) in a tissue-grinder for 40 min on ice, then centrifuged for 10 min at 15,000 *g* and 4°C. We used the Pierce BCA Protein Assay Kit (ThermoFisher) and a plate reader (Molecular Devices) to standardize protein loading in Laemmli 2X Concentrate sample buffer (Sigma) and deionized water for gel electrophoresis (Bio-Rad). Samples were heated to 95°C for 5 min on a heating block (Eppendorf) prior to loading. Membranes were probed with antibodies as above and imaged (LI-COR). We quantified band densities using Image Studio (LI-COR, v.5.2).

### Immunofluorescence microscopy

By our reported methods (26), we cannulated the trachea and pulmonary artery and incised the left ventricle of the heart of exsanguinated mice. We excised and inflated the lungs to 10 cmH_2_O, then perfused the lungs with 3 mL of ice-cold DPBS containing Ca^2+^ and Mg^2+^ followed by 5 mL of 4% methanol-free formaldehyde (ThermoFisher) in DPBS containing Ca^2+^ and Mg^2+^. The lungs were immersed in 4% methanol-free formaldehyde solution for up to 24 h, then rinsed and stored in 70% ethanol solution. Paraffin embedding, slicing (5 um), mounting, automated staining (Roche Diagnostics), and sealing was performed by the Pathology Department at the Icahn School of Medicine at Mount Sinai. Prior to staining, slides were immersed and heated for 1 min in Signal Stain Citrate Unmasking Solution (Cell Signaling), blocked for 30 min at room temperature with 5% goat serum (Jackson), and incubated for 40 min at room temperature with goat anti-mouse Fab fragments (Jackson). For imaging, we viewed slides by confocal microscopy (LSM800; Zeiss) using a 20x water immersion objective (NA 1.0; Zeiss) and randomly selected six regions of approximately 30-50 alveoli per lung for imaging. We confirmed absence of bleed-through between channels and acquired all images using identical microscope settings. Fluorescence was quantified using ImageJ (NIH; v.2.0.0-rc-69/1.52n) as mean intensity per image after thresholding to exclude airspaces that lacked fluorescence. Brightness and contrast adjustments were applied to individual color channels of entire images and equally to all experiment groups. We did not apply further downstream processing or averaging.

### Plasmid preparation, transfection, and instillation

We transformed DH5α *E. coli* (New England Biolabs) with plasmid DNA via heat-shock, then amplified and purified the plasmids using an EndoFree Plasmid Maxi Kit (Qiagen). By our established methods (26), we complexed plasmid DNA for A1440X (gift of Dr. Martina Gentzsch, University of North Carolina at Chapel Hill) or vector (pcDNA 3.1; Invitrogen) with freshy extruded unilamellar liposomes (20 ug/uL; 100 nm pore size; DOTAP; Avanti Lipids) in sterile Opti-MEM (Gibco). For transfection, we administered 75 ug plasmid DNA per mouse by intranasal instillation. We have shown previously (26) that this approach induces plasmid expression by the alveolar epithelium.

### Survival assessment

Mice were treated with instillations and injections as indicated in the figures. A blinded investigator assessed and recorded mouse weight, breathing score, and need for euthanasia in line with our IACUC-approved protocol. Need for euthanasia was determined by a scoring system that included observations of mouse appearance, breathing difficulty, grooming and eating behavior, gait, and response to stimulation by cage top opening and placement on a narrow beam. Mice instilled with IAV alone were assessed at 24 h intervals after IAV instillation for four days. Mice instilled with SA^GFP^ were assessed hourly for 6 h post-instillation, then at least every 12-24 h for 3 days. Surviving mice were euthanized at the conclusion of experiments.

### Protein and leukocyte determinations

Using our reported methods (26), we cannulated the trachea of exsanguinated mice, then lavaged the lungs with 5 sequential instillations of 1 mL of ice-cold, Ca^2+^-free PBS. For total protein determinations, we centrifuged the first 1 mL of BAL fluid for 10 min at 400 *g* and 4°C, then centrifuged the supernatant again for 20 min at 15,000 *g* and 4°C. Total protein was quantified using the Pierce BCA Protein Assay Kit (ThermoFisher) and a plate reader (Molecular Devices). For leukocyte determinations, BAL fluid samples were pooled on a per-mouse basis and centrifuged for 10 min at 500 *g*. The resuspended cells were incubated for 10 min in Turk’s solution (Sigma), then counted using a hemacytometer (Hausser Scientific).

### Virus and bacterial quantifications

We excised the lungs from exsanguinated mice using our reported methods (26). For bacterial quantification, lungs were mechanically homogenized by crushing in a specimen bag and diluted in 1 mL of DPBS containing Ca^2+^ and Mg^2+^. SA^GFP^ CFU was quantified by serial dilutions on chloramphenicol-containing LB agar plates. For viral quantification, lungs were homogenized in homogenizer tubes (Benchmark Scientific) in 500 uL of Ca^2+^-free PBS. IAV PFU was determined by plaque assay as described previously (72). Briefly, homogenized lungs were 10-fold serially diluted starting from 1:10 dilution and added to a confluent monolayer of Madin-Darby canine kidney cells (ATCC; line CCL-34) for 1 h at 37°C and 5% CO_2_ with gentle rocking. The inoculum was removed, and the cells were overlaid with a solution composed of 1% agar (Oxoid) and 2X minimal essential medium (MEM) supplemented with 1% diethyl-aminoethyl (DEAE)-dextran, 5% NaHCO_3_, and 1 ug/mL tosylamide-2-phenylethyl chloromethyl ketone (TPCK)-treated trypsin. Cell-containing plates were incubated for 48 h at 37°C and 5% CO_2_, then fixed in 10% formaldehyde overnight. Viral plaques were visualized by immune-labeling with mAb against HT-103 (gift of Thomas Moran’s laboratory, Icahn School of Medicine at Mount Sinai), HRP-conjugated anti-mouse secondary detection antibody, and TrueBlue substrate (KPL-Seracare).

### Statistics

Statistics are indicated in figures and legends. In general, paired comparisons were analyzed using two-tailed *t* tests, multiple comparisons were made using ANOVA with *post-hoc* testing, and survival comparisons were determined by log rank testing. We considered statistical significance at *P* < 0.05. Data were analyzed and figures were prepared using Microsoft Excel, StatPlus:mac Pro (AnalystSoft, Inc., version v7), and SigmaPlot (Systat, version 14.5).

### Study approval

The Institutional Animal Care and Use Committees of the Icahn School of Medicine at Mount Sinai and Columbia University Medical Center approved the animal procedures.

## AUTHOR CONTRIBUTIONS

J.H. designed the study, wrote the manuscript, and was responsible for the overall project. J.H., S.T., A.C.D., D.C., S.S., S.M., and R.R. contributed to data collection and data analysis. J.H., S.T., A.C.D., D.C., S.S., M.S., and J.B. contributed to the experimental design and interpretation of results. All authors edited the manuscript. The order of the co-first authors was determined based on S.T.’s greater contribution to the writing and composition of the figures.

## ACKNOWLEDGMENTS

This work was supported by grant HL164821, an American Heart Association Fellow-to-Faculty Transition Award 16FTF29380008, and a Columbia University Irving Institute/Clinical Trials Office Pilot Award (funded by UL1 TR001873) to J.H.; grants HL036024 and HL122730 to J.B.; and grant R21AI151229 to M.S. M. Garcia-Barros and R. Brody (Biorepository and Pathology CoRE, Icahn School of Medicine at Mount Sinai) provided assistance with immunofluorescence staining, and the laboratories of Drs. J. Bubeck-Wardenburg, M. Gentzsch, and T. Moran provided purified Hla, A1440X plasmid, and mAb against HT-103, respectively.

## REFERENCES

1. GBD 2017 Causes of Death Collaborators. Global, regional, and national age-sex-specific mortality for 282 causes of death in 195 countries and territories, 1980-2017: a systematic analysis for the Global Burden of Disease Study 2017. Lancet. 2018;392(10159):1736–88.

2. Jain S, et al. Community-acquired pneumonia requiring hospitalization among U.S. children. N Engl J Med. 2015;372(9):835–45.

3. Jain S, et al. Community-acquired pneumonia requiring hospitalization among U.S. adults. N Engl J Med. 2015;373(5):415–27.

4. Bartley PS, et al. Bacterial coinfection in influenza pneumonia: rates, pathogens, and outcomes. Infect Control Hosp Epidemiol. 2021:1–6.

5. Teng F, et al. Community-acquired bacterial co-infection predicts severity and mortality in influenza-associated pneumonia admitted patients. J Infect Chemother. 2019;25(2):129–36.

6. Rice TW, et al. Critical illness from 2009 pandemic influenza A virus and bacterial coinfection in the United States. Crit Care Med. 2012;40(5):1487–98.

7. Morens DM, et al. Predominant role of bacterial pneumonia as a cause of death in pandemic influenza: implications for pandemic influenza preparedness. J Infect Dis. 2008;198(7):962–70.

8. Baghdadi JD, et al. Antibiotic use and bacterial infection among inpatients in the first wave of COVID-19: a retrospective cohort study of 64,691 patients. Antimicrob Agents Chemother. 2021;65(11):e0134121.

9. Adalbert JR, et al. Clinical outcomes in patients co-infected with COVID-19 and *Staphylococcus aureus*: a scoping review. BMC Infect Dis. 2021;21(1):985.

10. Nickol ME, et al. Characterization of host and bacterial contributions to lung barrier dysfunction following co-infection with 2009 pandemic influenza and methicillin resistant *Staphylococcus aureus*. Viruses. 2019;11(2).

11. Finelli L, et al. Influenza-associated pediatric mortality in the United States: increase of *Staphylococcus aureus* coinfection. Pediatrics. 2008;122(4):805–11.

12. Bhattacharya J and Matthay MA. Regulation and repair of the alveolar-capillary barrier in acute lung injury. Annu Rev Physiol. 2013;75:593–615.

13. Huppert LA, et al. Pathogenesis of acute respiratory distress syndrome. Semin Respir Crit Care Med. 2019;40(1):31–9.

14. Randolph AG, et al. Vancomycin monotherapy may be insufficient to treat methicillin-resistant *Staphylococcus aureus* coinfection in children with influenza-related critical illness. Clin Infect Dis. 2019;68(3):365–72.

15. Guo Y, et al. Prevalence and therapies of antibiotic-resistance in *Staphylococcus aureus*. Front Cell Infect Microbiol. 2020;10:107.

16. Hussain M, et al. Drug resistance in influenza A virus: the epidemiology and management. Infect Drug Resist. 2017;10:121–34.

17. Plotkowski MC, et al. Adherence of type I *Streptococcus pneumoniae* to tracheal epithelium of mice infected with influenza A/PR8 virus. Am Rev Respir Dis. 1986;134(5):1040–4.

18. Pittet LA, et al. Influenza virus infection decreases tracheal mucociliary velocity and clearance of *Streptococcus pneumoniae*. Am J Respir Cell Mol Biol. 2010;42(4):450–60.

19. Sun K and Metzger DW. Influenza infection suppresses NADPH oxidase-dependent phagocytic bacterial clearance and enhances susceptibility to secondary methicillin-resistant *Staphylococcus aureus* infection. J Immunol. 2014;192(7):3301–7.

20. Martinez-Colon GJ, et al. Influenza-induced immune suppression to methicillin-resistant *Staphylococcus aureus* is mediated by TLR9. PLoS Pathog. 2019;15(1):e1007560.

21. Iverson AR, et al. Influenza virus primes mice for pneumonia from *Staphylococcus aureus*. J Infect Dis. 2011;203(6):880–8.

22. Sun K, et al. Nox2-derived oxidative stress results in inefficacy of antibiotics against post-influenza *S. aureus* pneumonia. J Exp Med. 2016;213(9):1851–64.

23. Wang C, et al. Influenza-induced priming and leak of human lung microvascular endothelium upon exposure to *Staphylococcus aureus*. Am J Respir Cell Mol Biol. 2015;53(4):459–70.

24. Stone KC, et al. Allometric relationships of cell numbers and size in the mammalian lung. Am J Respir Cell Mol Biol. 1992;6(2):235–43.

25. Dobbs LG and Johnson MD. Alveolar epithelial transport in the adult lung. Respir Physiol Neurobiol. 2007;159(3):283–300.

26. Hook JL, et al. Disruption of staphylococcal aggregation protects against lethal lung injury. J Clin Invest. 2018;128(3):1074–86.

27. Hageman JC, et al. Severe community-acquired pneumonia due to Staphylococcus aureus, 2003-04 influenza season. Emerg Infect Dis. 2006;12(6):894–9.

28. Londino JD, et al. Influenza matrix protein 2 alters CFTR expression and function through its ion channel activity. Am J Physiol Lung Cell Mol Physiol. 2013;304(9):L582–92.

29. Londino JD, et al. Influenza virus M2 targets cystic fibrosis transmembrane conductance regulator for lysosomal degradation during viral infection. FASEB J. 2015;29(7):2712–25.

30. Brand JD, et al. Influenza-mediated reduction of lung epithelial ion channel activity leads to dysregulated pulmonary fluid homeostasis. JCI Insight. 2018;3(20).

31. Lindert J, et al. Chloride-dependent secretion of alveolar wall liquid determined by optical-sectioning microscopy. Am J Respir Cell Mol Biol. 2007;36(6):688–96.

32. Wang PM, et al. Rapid alveolar liquid removal by a novel convective mechanism. Am J Physiol Lung Cell Mol Physiol. 2001;281(6):L1327–34.

33. Hummler E, et al. Early death due to defective neonatal lung liquid clearance in alpha-ENaC-deficient mice. Nat Genet. 1996;12(3):325–8.

34. Li T and Folkesson HG. RNA interference for alpha-ENaC inhibits rat lung fluid absorption in vivo. Am J Physiol Lung Cell Mol Physiol. 2006;290(4):L649–L60.

35. Borok Z, et al. Na transport proteins are expressed by rat alveolar epithelial type I cells. Am J Physiol Lung Cell Mol Physiol. 2002;282(4):L599–608.

36. Helms MN, et al. Dopamine activates amiloride-sensitive sodium channels in alveolar type I cells in lung slice preparations. Am J Physiol Lung Cell Mol Physiol. 2006;291(4):L610–8.

37. Cheek JM, et al. Tight monolayers of rat alveolar epithelial cells: bioelectric properties and active sodium transport. Am J Physiol. 1989;256(3 Pt 1):C688–93.

38. Cui G and McCarty NA. Murine and human CFTR exhibit different sensitivities to CFTR potentiators. Am J Physiol Lung Cell Mol Physiol. 2015;309(7):L687–99.

39. Kunzelmann K, et al. Mechanisms of the inhibition of epithelial Na(+) channels by CFTR and purinergic stimulation. Kidney Int. 2001;60(2):455–61.

40. Hegedus T, et al. Role of individual R domain phosphorylation sites in CFTR regulation by protein kinase A. Biochim Biophys Acta. 2009;1788(6):1341–9.

41. Alzamora R, et al. CFTR regulation by phosphorylation. Methods Mol Biol. 2011;741:471–88.

42. Prince LS, et al. Efficient endocytosis of the cystic fibrosis transmembrane conductance regulator requires a tyrosine-based signal. J Biol Chem. 1999;274(6):3602–9.

43. Parker D and Prince A. Immunopathogenesis of *Staphylococcus aureus* pulmonary infection. Semin Immunopathol. 2012;34(2):281–97.

44. Wang L, et al. Bacterial growth, detachment and cell size control on polyethylene terephthalate surfaces. Sci Rep. 2015;5:15159.

45. Zhang P, et al. Innate immunity and pulmonary host defense. Immunol Rev. 2000;173:39–51.

46. Westphalen K, et al. Sessile alveolar macrophages communicate with alveolar epithelium to modulate immunity. Nature. 2014;506(7489):503-6.

47. Zaynagetdinov R, et al. Identification of myeloid cell subsets in murine lungs using flow cytometry. Am J Respir Cell Mol Biol. 2013;49(2):180–9.

48. Su M, et al. Cytolytic peptides induce biphasic permeability changes in mammalian cell membranes. J Immunol Methods. 2001;252(1-2):63–71.

49. Lee MH, et al. A postinfluenza model of *Staphylococcus aureus* pneumonia. J Infect Dis. 2010;201(4):508–15.

50. Reznikov LR, et al. Antibacterial properties of the CFTR potentiator ivacaftor. J Cyst Fibros. 2014;13(5):515–9.

51. Bratcher PE, et al. Alterations in blood leukocytes of G551D-bearing cystic fibrosis patients undergoing treatment with ivacaftor. J Cyst Fibros. 2016;15(1):67–73.

52. Kong E and Jabra-Rizk MA. The great escape: pathogen versus host. PLoS Pathog. 2015;11(3):e1004661.

53. Engelhardt JF, et al. Expression of the cystic fibrosis gene in adult human lung. J Clin Invest. 1994;93(2):737–49.

54. Johnson MD, et al. Functional ion channels in pulmonary alveolar type I cells support a role for type I cells in lung ion transport. Proc Natl Acad Sci U S A. 2006;103(13):4964–9.

55. Brochiero E, et al. Evidence of a functional CFTR Cl(-) channel in adult alveolar epithelial cells. Am J Physiol Lung Cell Mol Physiol. 2004;287(2):L382–92.

56. Bove PF, et al. Human alveolar type II cells secrete and absorb liquid in response to local nucleotide signaling. J Biol Chem. 2010;285(45):34939–49.

57. Fang X, et al. Contribution of CFTR to apical-basolateral fluid transport in cultured human alveolar epithelial type II cells. Am J Physiol Lung Cell Mol Physiol. 2006;290(2):L242–9.

58. Regnier A, et al. Expression of cystic fibrosis transmembrane conductance regulator in the human distal lung. Hum Pathol. 2008;39(3):368–76.

59. Jiang J, et al. Pleural surface fluorescence measurement of Na+ and Cl-transport across the air space-capillary barrier. J Appl Physiol *(1985)*. 2003;94(1):343–52.

60. Solymosi EA, et al. Chloride transport-driven alveolar fluid secretion is a major contributor to cardiogenic lung edema. Proc Natl Acad Sci U S A. 2013;110(25):E2308–16.

61. Li X, et al. CFTR is required for maximal transepithelial liquid transport in pig alveolar epithelia. Am J Physiol Lung Cell Mol Physiol. 2012;303(2):L152–60.

62. Cui G, et al. VX-770-mediated potentiation of numerous human CFTR disease mutants is influenced by phosphorylation level. Sci Rep. 2019;9(1):13460.

63. Madden DR and Swiatecka-Urban A. Tissue-specific control of CFTR endocytosis by Dab2: Cargo recruitment as a therapeutic target. Commun Integr Biol. 2012;5(5):473–6.

64. Ameen N, et al. Endocytic trafficking of CFTR in health and disease. J Cyst Fibros. 2007;6(1):1–14.

65. van Riel D, et al. Human and avian influenza viruses target different cells in the lower respiratory tract of humans and other mammals. Am J Pathol. 2007;171(4):1215–23.

66. Ramos I, et al. Innate immune response to influenza virus at single-cell resolution in human epithelial cells revealed paracrine induction of interferon lambda 1. J Virol. 2019;93(20).

67. Peteranderl C, et al. Macrophage-epithelial paracrine crosstalk inhibits lung edema clearance during influenza infection. J Clin Invest. 2016;126(4):1566–80.

68. Takemura Y, et al. Cholinergic regulation of epithelial sodium channels in rat alveolar type 2 epithelial cells. Am J Physiol Lung Cell Mol Physiol. 2013;304(6):L428–37.

69. Chen XJ, et al. Influenza virus inhibits ENaC and lung fluid clearance. Am J Physiol Lung Cell Mol Physiol. 2004;287(2):L366–73.

70. Gavioli EM, et al. A current review of the safety of cystic fibrosis transmembrane conductance regulator modulators. J Clin Pharm Ther. 2021;46(2):286–94.

71. Kiefmann R, et al. Paracrine purinergic signaling determines lung endothelial nitric oxide production. Am J Physiol Lung Cell Mol Physiol. 2009;296(6):L901–10.

72. Jangra S, et al. Sterilizing immunity against SARS-CoV-2 infection in mice by a single-shot and lipid amphiphile imidazoquinoline TLR7/8 agonist-adjuvanted recombinant spike prote in vaccine. Angew Chem Int Ed Engl. 2021;60(17):9467–73.

