## Supplemental Figures 1-3 for "Rescue of alveolar wall liquid secretion blocks fatal lung injury by influenza-staphylococcal coinfection"

**Supplemental Figure 1**

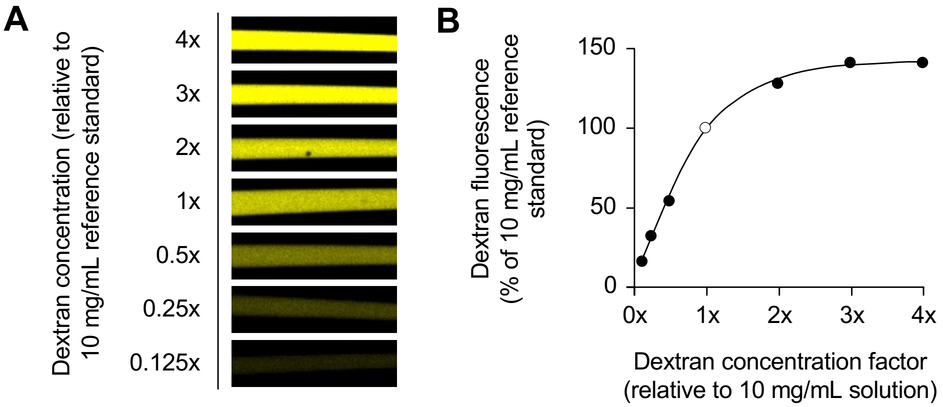


**Supplemental Figure 1. Dextran fluorescence calibration in glass micropipettes.** Confocal images (**A**) and plot (**B**) show the relationship between concentration and fluorescence intensity of tetramethylrhodamine (TRITC)-conjugated dextran (70 kD) in aqueous solution in glass micropipettes. *Open circle* (B) indicates the 10 mg/mL reference standard. *Line* calculated by polynomial regression ($P < 0.05$).

**Supplemental Figure 2**

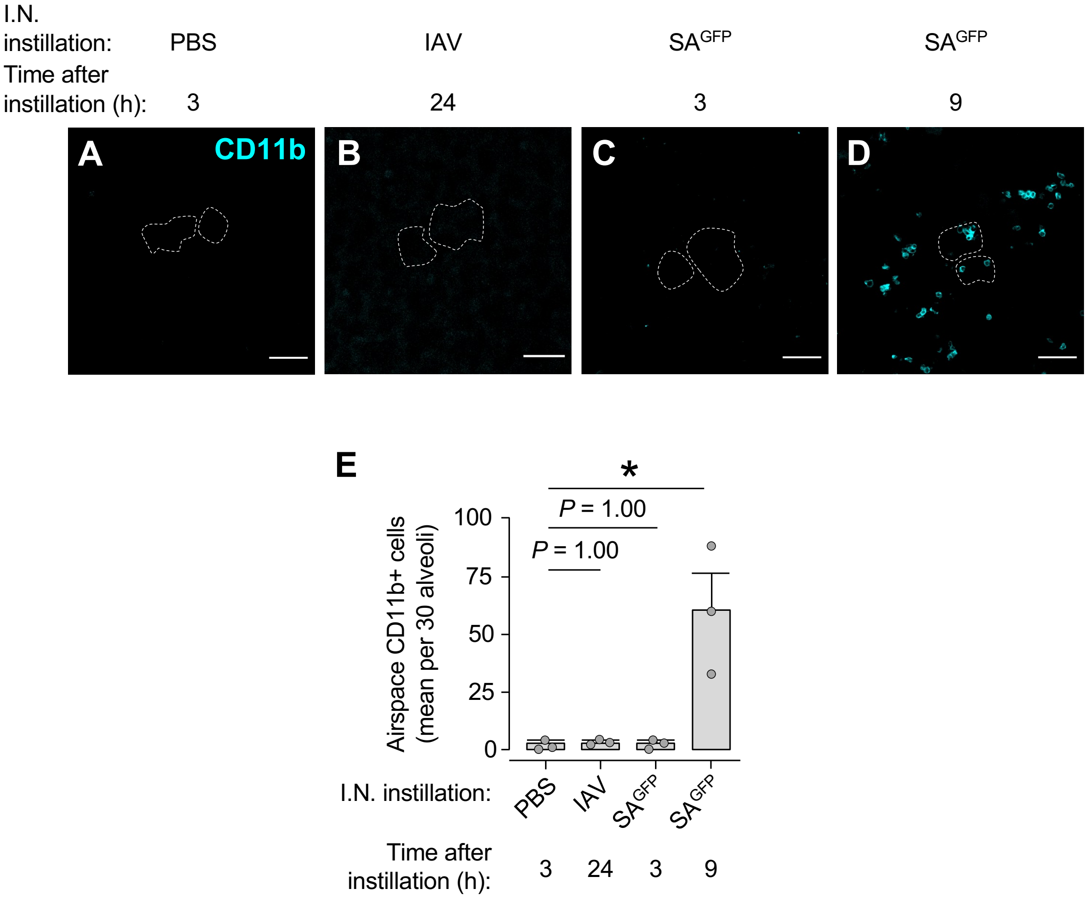


**Supplemental Figure 2. Leukocytes in airspaces of live alveoli after intranasal pathogen instillation.** Confocal images (**A-D**) and group data (**E**) show airspace fluorescence of allophycocyanin-conjugated CD11b antibody after alveolar antibody microinstillation in live, intact, blood-perfused mouse lungs. *Dotted lines* delineate example alveolar walls (fluorescence not shown). Imaging fields contain at least 30 alveoli each. Prior to imaging, mice were pretreated with intranasal instillation of PBS, IAV, or $SA^{GFP}$ as indicated, then the lungs were excised for imaging at the indicated time post-instillation. After antibody microinstillation, alveoli were microinstilled with HEPES-based buffer to remove non-specific antibody fluorescence. Note, CD11b fluorescence is apparent in alveolar airspaces only in lungs excised at 9 h after intranasal $SA^{GFP}$ instillation. For group data in E, circles indicate *n*, represent one mouse, and were generated by quantifying mean number of CD11b+ cells per imaging field of at least 30 alveoli. Bars: mean ± SEM; *$P < 0.05$ versus left bar by ANOVA with post hoc Tukey testing. Scale bars: 50 µm.

**Supplemental Figure 3**

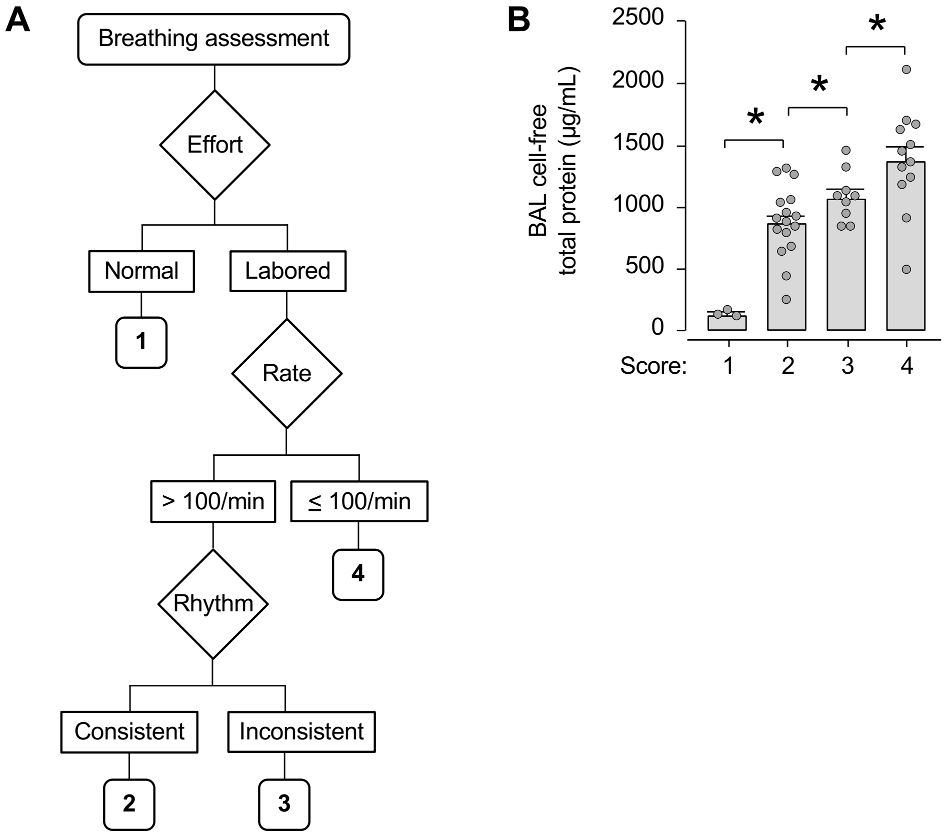


**Supplemental Figure 3. Breathing score assessment.** Investigators blinded to mouse group assigned scores to mice on a 1-4 scale using the flow diagram indicated in **A**. Group data in **B** show the correlation between breathing scores and total protein content in BAL fluid of pathogen-instilled mice. Circles (B) indicate *n* and each represent BAL protein content in one mouse. To generate the group data, we assigned breathing scores to mice infected with IAV, $SA^{GFP}$, or both, then plotted the relationship between score and protein content. Bars: mean ± SEM; *$P < 0.05$ by two-tailed *t* test as indicated.
